# Type-I interferon signaling is essential for robust metronomic chemo-immunogenic tumor regression in murine triple-negative breast cancer

**DOI:** 10.1101/2021.12.05.471293

**Authors:** Cameron Vergato, Kshama A. Doshi, Darren Roblyer, David J. Waxman

## Abstract

Triple-negative breast cancer (TNBC) is characterized by poor prognosis and aggressive growth, with limited therapeutic options for many patients. Here, we use two syngeneic mouse TNBC models, 4T1 and E0771, to investigate the chemo-immunogenic potential of cyclophosphamide and the mechanistic contributions of cyclophosphamide-activated type-I interferon (IFN) signaling to therapeutic activity.

Chemically-activated cyclophosphamide induced robust IFNα/β receptor-1-dependent signaling linked to hundreds of IFN-stimulated gene responses in both TNBC lines. Further, in 4T1 tumors, cyclophosphamide given on a medium-dose, 6-day intermittent metronomic schedule induced strong IFN signaling but comparatively weak immune cell infiltration associated with long-term tumor growth stasis. Induction of IFN signaling was somewhat weaker in E0771 tumors but was followed by extensive downstream gene responses, robust immune cell infiltration and prolonged tumor regression. The immune dependence of these effective anti-tumor responses was established by CD8 T-cell immunodepletion, which blocked cyclophosphamide-induced E0771 tumor regression and led to tumor stasis followed by regrowth. Strikingly, IFNα/β receptor-1 antibody blockade was even more effective in preventing E0771 immune cell infiltration and blocked the major tumor regression induced by cyclophosphamide treatment. Type-I IFN signaling is thus essential for the robust chemo-immunogenic response of these TNBC tumors to cyclophosphamide administered on a metronomic schedule.

**Significance:** TNBC has poor prognosis and few therapeutic options. We show that cyclophosphamide treatment can induces extensive tumor regression in syngeneic mouse models of TNBC via a chemo-immunogenic mechanism linked to type-I IFN production. Our findings establish that IFN signaling is essential for the robust anti-tumor actions of cyclophosphamide and suggest that treatment resistance may stem from silencing the IFN pathway. This suggests a new avenue for improving TNBC treatment efficacy.

## Introduction

Triple-negative breast cancer (TNBC) is characterized by increased tumor aggression and poor prognosis compared to other breast cancer subtypes (1). TNBC is distinguished by the lack of estrogen receptor, progesterone receptor and human epidermal growth factor receptor 2 (HER2) (2), which limits treatment options. TNBC initially responds to neoadjuvant chemotherapy but often recurs and metastasizes, with poor patient prognosis. Checkpoint inhibitors are often ineffective in TNBC patients (3,4), despite a comparatively high mutational burden and elevated levels of tumor-infiltrating lymphocytes (5). The discovery and preclinical development of novel therapies is thus critically important.

Immunogenic cell death is a unique cell death mechanism that can activate both innate and adaptive immune responses (6,7) and confer long term immunity (8). Chemotherapy-induced immunogenic cell death is characterized by damage-associated molecular pattern responses (9), including cell surface translocation of calreticulin, a pro-phagocytic signal (10), release of the toll-like receptor 4 ligand HMGB1 (11), extracellular release of ATP (12) and production of type-I interferons (IFN) (13). Dendritic cells attracted by the release of these molecules by dying tumor cells in the tumor microenvironment engulf the dead and dying tumor cells and undergo maturation (7). Immunostimulatory cytokines produced by mature dendritic cells, in turn, recruit NK cells, CD4 T-cells and CD8 T-cells, which can contribute to tumor regression and activate tumor-specific immunity (7). Several cytotoxic drugs approved for breast cancer have the potential to induce immunogenic cell death, including doxorubicin (14), epirubicin (15), mitoxantrone (7) and cyclophosphamide (CPA) (16,17).

Type-I IFNs, primarily IFNα and IFNβ, are secreted in response to viral or bacterial infection when viral gene products or bacterial cell wall components are detected by toll-like receptors or by cytosolic sensors of specific nucleic acids (18). Type-I IFNs bind to the heterodimeric IFN α/β receptor (IFNAR), which in turn activates a signaling cascade leading to increased expression of many interferon-stimulated-genes (ISGs). These ISGs have diverse immunomodulatory effects, including immune cell recruitment, type-2 IFN production and immune cell activation (18), opening up many novel interferon-based therapeutic opportunities for cancer treatment (19). Type-I IFN signaling supports tumor cell immunosurveillance (20) and impacts the efficacy of certain anti-cancer therapies, including antibodies against HER2, anthracyclines, checkpoint inhibitors, and lenalidomide (21).

In murine glioma models, CPA can induce immunogenic cell death when administered on a metronomic, medium-dose intermittent chemotherapy (MEDIC) schedule (22,23), leading to elimination of GL261 gliomas implanted in syngeneic mice and activation of long-term anti-tumor immunity (24). Other CPA treatment schedules are much less effective at inducing immune cell recruitment in glioma models (25), a finding that was recently validated in breast cancer models (26). CPA given on a MEDIC schedule activates tumor cell autonomous type-I IFN signaling required for CPA-induced immune cell infiltration (17), suggesting cytotoxic drug-induced type-I IFN production may serve as a biomarker for the immunogenic potential of cancer cells. However, it is not known whether, and to what extent, IFN-stimulated immune cell recruitment contributes to the tumor regression induced by MEDIC CPA treatment.

Here, we investigate the immunogenic potential of CPA in two TNBC tumor models: 4T1, a Balb/c mouse syngeneic mammary carcinoma model for metastatic late-stage breast cancer (27); and E0771, a medullary breast adenocarcinoma formed spontaneously in C57BL/6 mice (28) and model for spontaneous breast cancer (29). Orthotopic E0771 tumors undergo CD8 T-cell-dependent tumor regression with specific anti-tumor immunity when treated with doxorubicin combined with interleukin-2 (30), but immune-based tumor regression induced by chemotherapy alone, including MEDIC scheduling of CPA (22,23), has not been reported for either TNBC model.

We assay these TNBC lines for their capacity to mount an interferon response, as indicated by rbust interferon-stimulated-gene (ISGs) induction following *in vitro* treatment with 4HC or doxorubicin, and we assess the dependence on IFNα/β receptor-1 (IFNAR-1) signaling. Further, we investigate the impact of CPA administered on a MEDIC metronomic schedule on TNBC tumors implanted orthotopically in syngeneic mice. Our findings reveal a striking immunogenic response to CPA associated with increased expression of hundreds of genes, including many ISGs, and resulting in the near complete regression of E0771 tumors in a manner that is absolutely dependent on the activation of type-I IFN signaling-supported immune cell recruitment.

## Materials and Methods

### Tumor cell lines

Cell lines were authenticated by and obtained from CH3 BioSystems (Amherst, NY) (E0771, cat. #94A001) and American Type Culture Collection (Manassas, VA) (4T1 cells, cat. #CRL-2539; B16F10 cells, cat. #CRL-6475). Typically, cell lines were propagated in culture for fewer than 6-8 passages before cells were discarded and a fresh, early passage cell vial was thawed and used for experimentation. Cells were cultured in RPMI-1640 (4T1, EO771) or DMEM (B16F10) medium, 10% fetal bovine serum (FBS) and 1% penicillin-streptomycin at 37°C under a humidified 5% CO_2_ atmosphere. Cells were stained with 0.4% trypan blue and counted using a Countess Automated Cell Counter (Thermo-Fisher Scientific).

### Cytotoxicity/chemosensitivity (MTS) assa

Cells were seeded in 96-well plates, (cat. #10861-666, VWR, Radnor, PA) at 3,000 cells per well, 1 day prior to treatment with 10^−9^ M to 10^−4^ M chemically activated CPA (4-hydroperoxy-CPA, 4HC; cat. # 19527, Cayman Chemical, Ann Arbor, MI) or doxorubicin (cat. #D1515 Sigma-Aldrich) for 4 h. Cells were then washed once with PBS (cat. #BP24384, Fisher Scientific), cultured for 68 h in drug-free media. MTS reagent (10 µl; cat. # G5421, Promega, Madison, WI) then incubated at 37°C to assay cell viability. A_490_ was measured every 30 min (Synergy H1 plate reader; BioTek Instruments, Winooski, VT), and the time-point where untreated cells reached A_490_= 1.0 was used to generate dose-response viability curves and calculate IC_50_ values by non-linear curve fitting implemented in GraphPad Prism 8.

### In vitro drug treatment

Cells were treated for 4 h with 4HC or doxorubicin at IC_50_-range drug concentrations using cells seeded the prior day at 50,000 (E0771) or 75,000 cells per well (4T1, B16F10) of a 6-well plate (cat. #10861-696, VWR). Cells were then washed once with PBS and incubated in fresh media for a total of 24, 48 and 72 h after the start of drug treatment, at which time RNA was isolated.

### RNA isolation and quantitative PCR (qPCR)

RIzo™Reagent (1 ml; cat. # 15596018, Invitrogen, Carlsbad, CA) was used to extract RNA from ∼30-200 mg frozen tumor tissue or from cells in one well of a 6-well plate. RNA was resuspended in ultrapure water and quantified (BioTek Synergy H1 plate reader or Qubit(tm) 3.0 Fluorometer) (cat. #15387293, Fisher Scientific). RNA (1 µg) was treated with RNase-free RQ1 DNase 1 (cat. # M6101, Promega) with a murine RNase inhibitor (cat. #M0314, New England Biolabs) followed by cDNA synthesis using a High-Capacity cDNA Reverse Transcription kit (cat. # 466814, Applied Biosystems, Foster City, CA). qPCR was performed on cDNA samples using *Power* SYBR™ Green PCR Master Mix (cat. # 4367659, Applied Biosystems), gene-specific primers (Table S1) (Eton Bioscience, San Diego, CA) and a BioRad CFX384 Touch™ Real-Time PCR Detection System.

Data for mouse ISGs (Mx1, Cxcl10, Oasl1, Cxcl11, Igtp, RSAD2) and immune marker genes (Cd8α, Nkp46, Cd68, Ifng, Prf1, Gzmb, Cd11b and Foxp3) was analyzed by the comparative Ct method. Gene expression, normalized to 18S RNA content, was presented relative to untreated cells for *in vitro* samples, or to placebo group for *in vivo* tumor samples. Target gene primers pairs were designed to span two adjacent exons, to be 18-22 bp long with close to 50% G:C content, and to form amplicons 50-150 bp long. Unique primer specificity was verified by extending each primer sequence by 3, 5, 10, 15 and 20 nucleotides and then using the UCSC Genome Browser BLAST-like alignment tool (BLAT) to confirm a single correct target. Data for culture experiments is presented as mean +/-standard deviation (SD) with n = 2-3 replicate samples. Mouse experiments are presented as mean +/-standard error of the mean (SEM) for n tumors, as indicated. qPCR primer sequences are shown in Fig. S8.

### In vitro interferon-β (IFNβ) treatment

Cells seeded in 6-well plates at 200,000 cells per well were incubated overnight, then treated with recombinant mouse IFNβ1 (cat. # 581302, BioLegend, San Diego, CA) at 28, 83 or 250 U/mL for 4 h. Cells were then washed with PBS, fresh media was replaced, and cells were harvested 2 h later for RNA isolation.

### poly (I:C) transfection

Cells were transfected with 1 µg/mL poly (I:C) (cat. # tlrl-picw, InVivogen, San Diego, CA) using 6 µg/mL poly-ethylenimine. Cells were seeded overnight in 6-well plates at 50,000 cells/well for E0771 cells and 75,000 cells/well for 4T1 and B16F10 cells. The next day, poly (I:C) (2 µL of 1 mg/mL per well) was mixed with 12 µL of 1 mg/mL poly-ethylenimine and 100 µL of serum-free media and incubated at room temperature for 15 min. This solution was added to 1.89 mL of full media and placed in one well of a 6-well plate for 4 h. Cells were then washed with PBS, followed by addition of fresh media. Cells were collected 20 h later for RNA isolation.

### In vitro interferon receptor antibody treatment

Cells were treated with 10 µg/mL monoclonal anti-mouse IFNα/β receptor subunit 1 (IFNAR-1) antibody (clone MAR1-5A3, BioXCell, West Lebanon, NH), which was added to the cells together with 4HC, doxorubicin, IFNβ or poly (I:C), for 4 h, as above. The media was removed, and the cells were washed once in PBS before adding fresh media containing 10 µg/mL IFNAR-1 antibody for an additional 2 h to 70 h prior to harvesting for RNA isolation.

### Conditioned media treatment

4T1 and E0771 cells were treated with 4HC (5 µM and 4.2 µM, respectively) as described above and harvested 72 h later. Conditioned media was collected from these donor cells, transferred to drug-free (naïve) recipient cells seeded overnight in 6-well plates, and incubated for 4 h. Cells were washed with PBS followed by replacement with fresh media prior to isolation of RNA from the recipient cells 2 h later.

### Mouse studies: tumor inoculation and CPA treatment

Mice were treated using protocols specifically reviewed for ethics and approved by the Boston University Institutional Animal Care and Use Committee (protocol # PROTO201800698), and in compliance with ARRIVE 2.0 Essential 10 guidelines (31), including study design, sample size, randomization, experimental animals and procedures, and statistical methods. 6-week-old female BALB/c mice (Taconic Farms, Germantown, NY) and female C57/BL6N mice (Taconic Farms) were purchased as indicated. 4T1 cells (1 × 10^5^) or E0771 cells (2 × 10^5^) resuspended in 0.1 ml PBS were inoculated into the fourth mammary fat pad of BALB/c and C57/BL6N mice, respectively, using a 1 mL syringe (cat. #309628, BD Biosciences) with a 5/8-inch-long 26-gauge needle (cat. #305115, BD Biosciences, San Jose, CA). Tumor length and width were monitored every 3 d using a vernier caliper, and tumor volumes were calculated: Volume = (π/6) × (L × W)^3/2^. Mice were randomized into treatment and placebo groups when average volumes reached 100-150 mm^3^ (4T1) or 200-250 mm^3^ (E0771). CPA (cat. # C0768, Sigma-Aldrich, St. Louis, MO) dissolved in sterile PBS and passed through a 0.2 µm filter was then injected i.p. at 0 (vehicle control), 90, 110 or 130 mg/kg, as indicated. CPA injections were repeated every 6 days. Mice were euthanized at specified time points. Tumors were excised, washed with PBS and flash-frozen in liquid nitrogen after placing ∼1/3 piece of fresh tumor in 1 mL TRIzol for immediate downstream use, as required.

### Immunodepletion studies

To deplete CD8 T-cells, 0.28 mg of anti-mouse CD8α antibody (clone 53-6.7, cat. #BE0004-1, BioXCell) or rat IgG (cat. #I4131, Sigma-Aldrich), was diluted in 0.1 ml sterile PBS then given to mice by IP injection repeated on days -5, -1, 3, 9 and 15 (c.f., 110 mg/kg CPA treatment beginning on day 0). To achieve IFNAR-1 blockade, anti-mouse IFNAR-1 antibody or mouse IgG (cat. #MS-GF-ED, Molecular Innovations, Novi, MI) diluted in sterile PBS and injected i.p. (as above) as follows: 1.0 mg on day -1, 0.5 mg on day 0, and 0.25 mg on days 3, 6, 9 and 12, with 110 mg/kg CPA treatment every 6 days, beginning on day 0.

### Fluorescence-activated cell sorting (FACS) of tumor and blood samples

Approximately 1/3 of each freshly harvested tumor was dissociated to generate a 0.5 ml single-cell suspension using a Miltenyi Biotec gentleMACS™ Dissociator (cat. #130-093-235,), C-tubes (cat. #130-093-237) and Mouse Tumor Dissociation Kit (cat. #130-096-730) by using the manufacturer’s instructions for “tough” tumor samples. Mouse blood obtained by tail-vein blood collection (20 μL) was placed in a 1.5 mL microcentrifuge tube with 5 μL of 1000 U/mL heparin sodium (cat. #H3393, Sigma Aldrich) in 0.9% saline. 1 mL of 1X RBC Lysis Buffer (cat. #00-4333-57, Thermo-Fisher Scientific) was then added to 25 uL of each sample (dissociated tumor samples or blood samples) and shaken for 20 min at 20°C to destroy red blood cells. Cells were then washed with 2 mL PBS and centrifuged at 400 × g for 5 min, followed by a second wash with 3 mL Protein Extraction Buffer (PEB: 0.5% BSA, 2 mM EDTA in PBS, pH 7.2). The cells were spun at 400 × g spin for 5 min and resuspended in 200 μL of PEB. 100 μL was removed, mixed with 2 μL of anti-mouse CD16/CD32 antibody (cat. #14-0161-85, Thermo-Fisher Scientific), and incubated at 4°C for 20 min to block nonspecific IgG binding. Anti-mouse CD8α-APC antibody (0.7 μL; cat. #20-1886, Clone 2.43, Tonbo Biosciences, San Diego, CA) was added then incubated for 10 min at 4°C. Cells were then washed with 3 mL of PEB, spun for 5 min at 400 × g and resuspended in 200 μL PEB. Propidum iodide (cat. #P3566, Thermo-Fisher Scientific) was added (20 ng/mL, final concentration), followed immediately by processing on a BD Biosciences FACSCalibur instrument (cat. #342975) and analysis using BD CellQuest Pro Software (BD Biosciences). Counted events were first gated by size based on forward scattering and side scattering parameters to omit very large and very small events. The next gate separated living from dead cells by excluding events with a propidium iodide signal. CD8α+ cells were then counted by excluding events lacking an APC signal. CD8α+ cells were presented as a percentage of total live cells.

### RNA-seq library preparation and sequence analysis

Polyadenylated mRNA was isolated from 1 μg of total RNA from cultured tumor cells or excised tumor tissue using NEBNext^®^ Poly(A) mRNA Magnetic Isolation Module (cat. #E7490, New England Biolabs) and the manufacturer’s instructions. The resulting polyA-selected RNA was used to prepare RNA-seq libraries using the NEBNext^®^ Ultra^(tm)^ II Directional RNA Library Prep Kit for Illumina^®^ (cat. #E7600, New England Biolabs), NEBNext^®^ Multiplex Oligos for Illumina^®^ Dual Index Primers Set 1 (cat. #E7600, New England Biolabs), and AMPure^®^ XP Beads (cat. #A63881, Beckman Coulter Inc., Indianapolis, IN) per the manufacturer’s instructions. Cell culture-derived RNA-seq libraries were prepared for n=3 independent cultures for each condition tested, except for vehicle control-treated 4T1 cells, where n=2. Tumor-derived RNA-seq libraries were prepared for n=2-3 independent pools for tumor-extracted RNA for each condition tested, with each pool prepared from n=2-3 independent biological replicate tumors. Libraries were multiplexed and sequenced by Novogene, Inc (Sacramento, CA) to an average depth of 28 million (cell culture libraries) or 13 million (tumor-derived libraries) paired-end sequence reads each (Table S3). Data was analyzed using an in-house custom RNA-seq pipeline (32), using edgeR (33) to identify differentially expressed genes for each indicated comparison, using the following cutoffs: p-value < 0.05, |fold-change| > 2.0, and fragments per kilobase per million reads (FPKM) > 0.5-1.0, as specified. Gene lists were input into the DAVID Bioinformatics Database’s Functional Annotation Clustering Tool (https://david.abcc.ncifcrf.gov/home.jsp) to identify functional enrichment clusters with significance scores for each gene list. Data are presented for the top enriched term for each of the top three clusters, along with their Benjamini-Hochberg adjusted p-values.

### Data availability

The data generated in this study are available within the article and its supplementary data files. High throughput sequencing data (Fastq files and processed data files) are available for download from Gene Expression Omnibus (GEO) (https://www.ncbi.nlm.nih.gov/geo/) at accession # GSEXX.

## Results

### 4HC induces a type-I IFN response in breast cancer lines

4T1 and E0771 breast cancer cells were treated with 4HC, an activated CPA metabolite (34), or with doxorubicin, an established immunostimulatory chemotherapeutic drug (7,14). Drug exposures were for 4 h (t = 0 to 4 h) using IC_50_-range drug concentrations (Fig. S1) to mimic *in vivo* exposure to the activated CPA metabolite 4-hydroxy-CPA, which is >90% cleared from mouse circulation within 4-h of CPA dosing (35). Total cellular RNA was extracted and assayed for changes in the expression of several ISGs. 4HC induced strong increases in all three ISGs examined after a 24-48-h lag, with responses being stronger in E0771 cells than 4T1 cells (Fig. 1B vs. Fig. 1A). ISG responses to doxorubicin were stronger than responses to 4HC in E0771 cells but were weaker in 4T1 cells. ISG induction was also observed when culture supernatant from 4HC-treated 4T1 and E0771 cells was applied to drug-naïve cells, indicating that the drug-treated cells secrete ISG-stimulatory cytokines (Fig. S2), such as the type-I IFNs, IFNα and IFNβ. Weak ISG responses were seen in B16F10 melanoma cells treated with 4HC or doxorubicin at their IC_50_ concentrations (Fig. 1C; Fig. S1C). This finding is consistent with the weak immune responses seen in CPA-treated B16F10 tumors implanted in syngeneic mice (36).

**Fig. 1.**
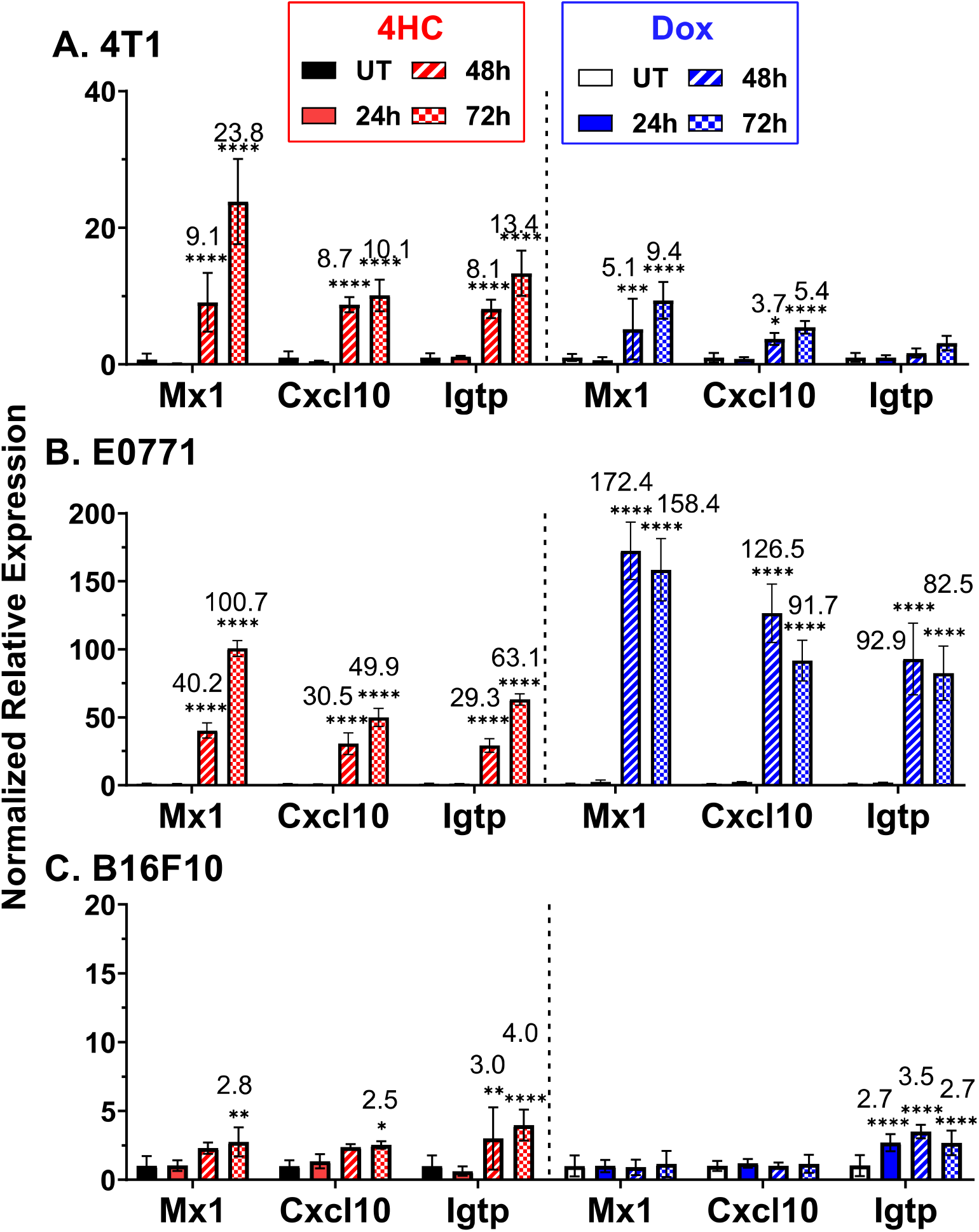
4HC and doxorubicin induction of ISGs in cultured tumor cell lines. **A**. 4T1 cells were treated for 4 h with IC_50_-range concentrations of 4HC (5 µM) or doxorubicin (2 µM) (see Fig. S1), followed by removal of drug and further incubation until 24, 48 or 72 h after initiating drug treatment. RNA was then extracted and analyzed by qPCR for expression of the three indicated ISGs. **B**. E0771 cells were treated with 4HC (4.2 µM) or doxorubicin (1.7 µM) as described in A. **C**. B16F10 cells treated with 4HC (20 µM) or doxorubicin (6.6 µM). Data shown are mean +/-SD for n = 2-3 replicates, with statistical significance determined by 2-way ANOVA implemented in GraphPad Prism: *, p < 0.05; **, p < 0.01; ***, p < 0.001; ****, p < 0.0001.

### ISGs are induced by multiple signaling pathways in 4HC-treated TNBC cells

To assess the role of type-I IFN signaling in these ISG responses, 4T1 and E0771 cells were treated with 4HC in combination with anti-IFNAR1 antibody under conditions that effectively blocks direct type-I IFN responses (Fig. S3). In both breast cancer cell lines, anti-IFNAR1 antibody completely blocked 4HC induction of *Igtp*, but only partially inhibited the induction of *Oasl1* and *Cxcl10* (Fig. 2A, Fig. 2B). Thus, IFNAR1 signaling contributes to, but does not entirely explain, the latter two ISG responses to 4HC treatment. To further investigate the underlying mechanism for ISG induction, cells were transfected with the ds-RNA analog, poly I:C, both with and without anti-IFNAR1 antibody. All three ISGs were induced by poly I:C in both cell lines, but only the 4T1 cell response was completely blocked by anti-IFNAR1 antibody (Fig. 2C, Fig. 2D). Thus, E0771 cells showed a pattern of partial inhibition of ISG induction by anti-IFNAR1 antibody with both 4HC and poly I:C. These findings are consistent with the proposal that 4HC activates a dsRNA-dependent mechanism leading to an increase in type-I IFN production and the observed downstream ISG responses. The partial inhibition by anti-IFNAR1 antibody of *Oasl1* and *Cxcl10* induction indicates these ISGs can also be formed by a type-I IFN-independent mechanism in 4HC-treated cells.

**Fig. 2.**
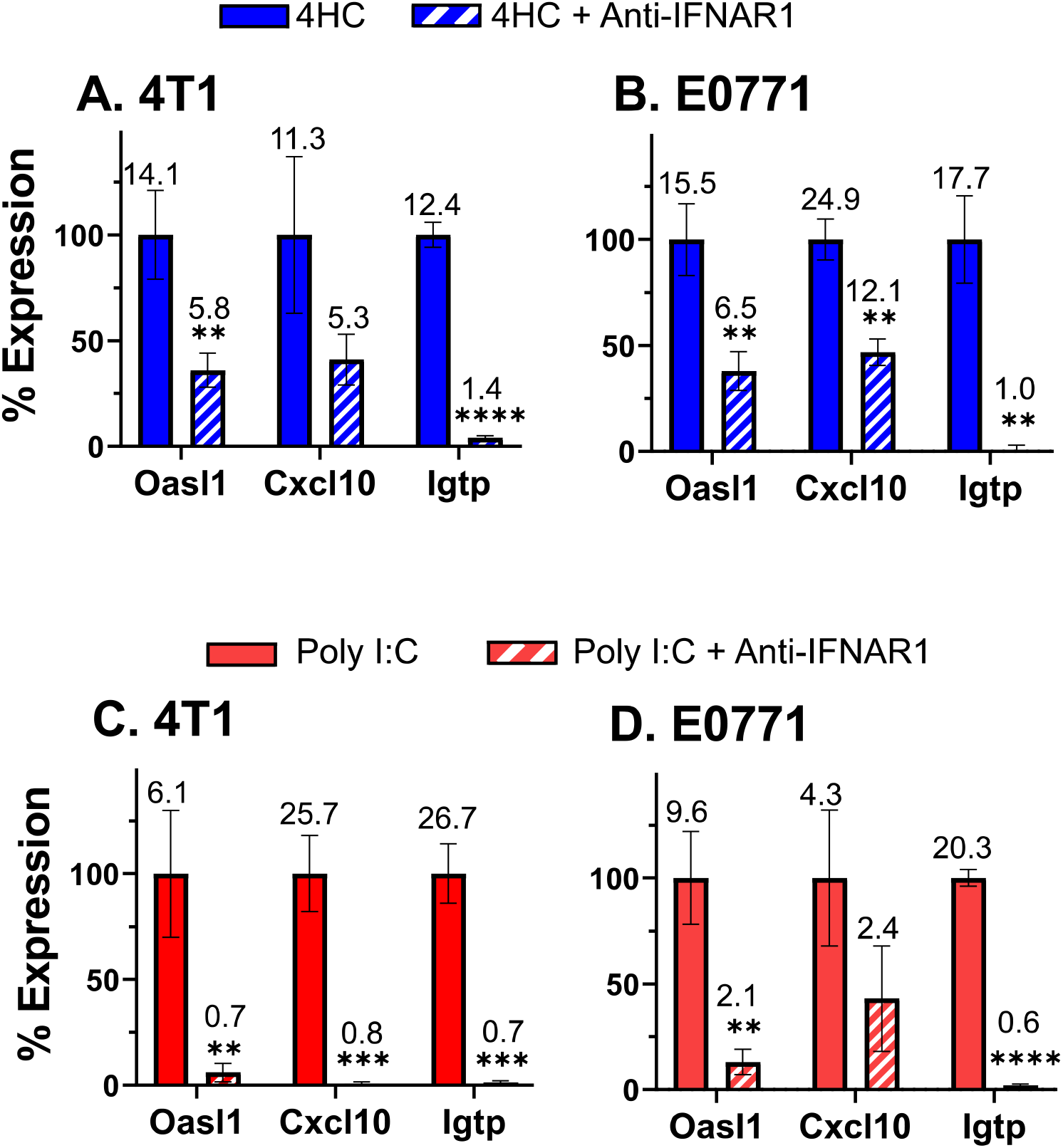
Anti-IFNAR-1 antibody Inhibits ISG induction by 4HC or poly (I:C) **A, B**. 4T1 and E0771 cells were treated with anti-IFNAR-1 antibody (10 μg/mL; Fig. S3) in combination with 4HC using the 72 h time-point protocol of Fig. 1, followed by qPCR analysis for ISG induction. **C, D**. 4T1 and E0771 cells were treated with 1 μg/mL poly (I:C) for 4 h, alone or in combination with anti-IFNAR-1 antibody (10 μg/mL), then further incubated for 20 h in the presence of IFNAR-1 antibody followed by qPCR analysis of ISG induction. Data presented are mean +/-SD values for n=2-3 replicates, with significance of the effect of antibody assessed by t-test: *, p < 0.05; **, p < 0.01; ***, p< 0.001; ****, p < 0.0001. Percent gene expression was calculated as: ((x ± SD) − (z ± SD))/((y ± SD) − (z ± SD)), where × = 4HC or poly (I:C) with anti-IFNAR-1 expression value, z = untreated control expression value, and y = 4HC or poly (I:C) alone expression value. Fold-change values are listed above each bar. Results shown are representative of at least 2 or 3 independent experiments.

### Global effects of 4HC exposure

RNA-seq was used to characterize the global impact of 4HC exposure on both type-I IFN-dependent and type-I IFN-independent genes that may potentially contribute to downstream immunostimulatory responses in each breast cancer line. In 4T1 cells, 4HC induced expression of 1043 genes, of which 388 (37%) also responded to short-term stimulation with recombinant IFNβ, which identifies the latter genes as 4T1 breast cancer type-I ISGs (Fig. 3A). Similarly, in E0771 cells, 188 (34%) of 568 genes induced by 4HC were also induced by IFNβ (Table S1). Top functional annotation clustering terms include innate immunity and virus response (Fig. 3A, Table S2), consistent with 4HC inducing many type-I IFN response genes in both breast cancer lines. Further supporting the proposed activation of type-I IFN signaling by 4HC in both TNBC models, we identified 110 genes induced by both 4HC and IFNβ in both cell lines, and for 97 of these genes, anti-IFNAR1 antibody significantly inhibited gene induction by 4HC (Table S1C). Finally, in 4T1 cells but not E0771 cells, many other genes were suppressed by 4HC, with enrichment for mRNA splicing and ribosome biogenesis. Notably, 85 of these genes were also suppressed by IFNβ treatment (Fig. 3B, Table S1, Table S2).

**Fig. 3.**
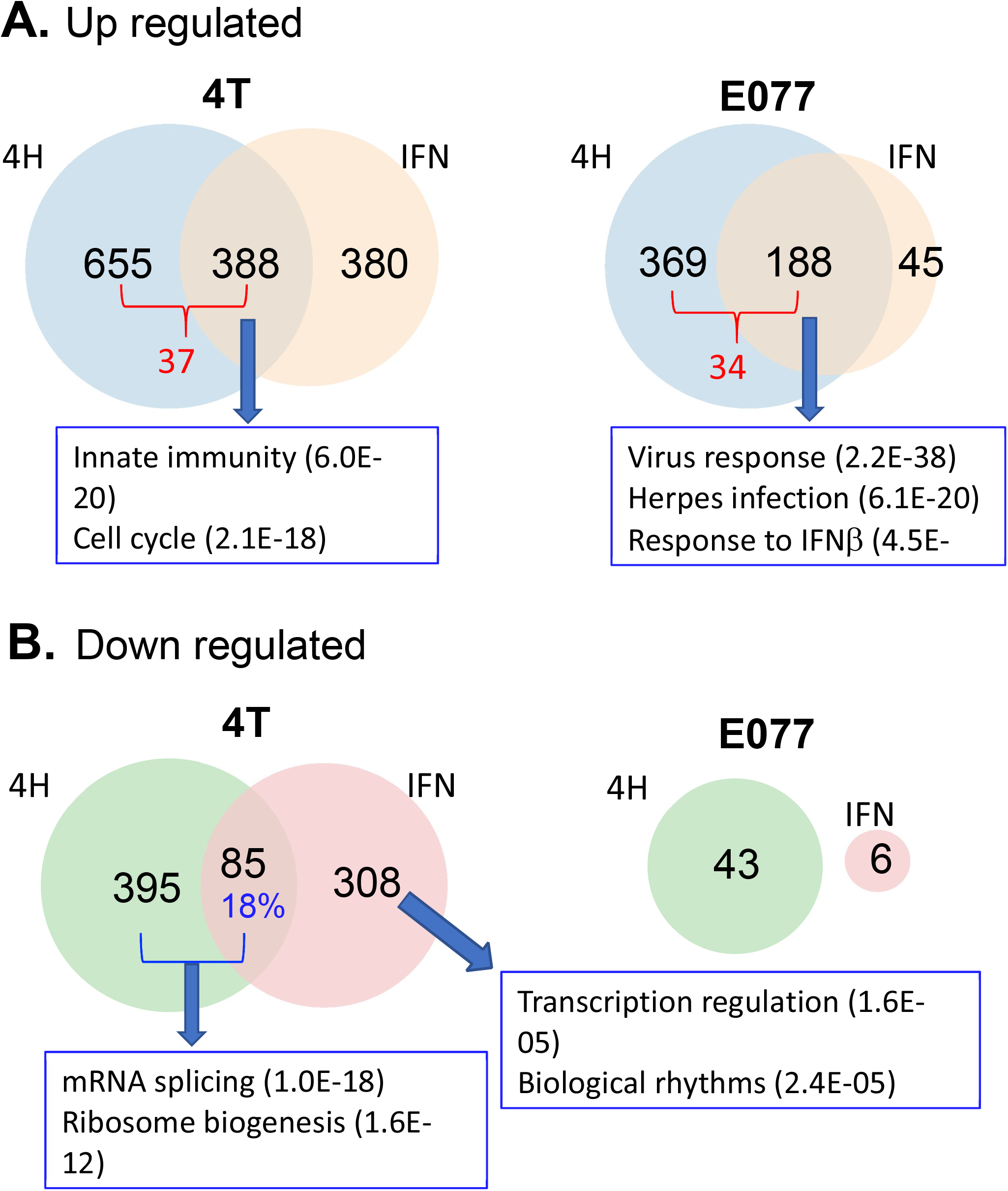
Gene responses to 4HC and IFNβ in cultured 4T1 and E0771 cells: RNA-seq analysis. Venn diagrams showing numbers of genes induced (**A**) or repressed (**B**) in cells treated with 4HC (4 h exposure, harvest 68 h later, as in Fig. 1) or recombinant IFNβ (4 h exposure, 2 h harvest 2 h later) based on a >2-fold change in expression at FDR < 0.05. Full gene lists are shown in Table S1. Top functional annotation clustering terms for the indicated gene sets, and their enrichment significance, are shown in boxes. The full DAVID analysis is presented in Table S2.

### Metronomic CPA induces 4T1 tumor growth stasis

We investigated the impact of CPA treatment on growth of implanted 4T1 tumors, ISG expression, and immune cell recruitment. Female BALB/c mice with orthotopic 4T1 tumors were treated with CPA (130 mg/kg) or placebo (PBS) on an intermittent metronomic, 6-day repeating schedule (MEDIC schedule; (22)). CPA dramatically reduced the rapid growth seen in drug-free tumors (placebo group) within one treatment cycle, inducing growth stasis that persisted through 7 cycles (Fig. 4A). When treatment was halted after 4 CPA cycles, tumor growth resumed 12 days later (Fig. 4A, day 36). Analysis of total tumor RNA extracted after 2, 4 and 7 CPA cycles revealed that the ISGs *Cxcl10* and *Mx1* were initially upregulated but returned to baseline after discontinuation of CPA treatment (Fig. 4B). We evaluated tumor immune cell infiltration by monitoring changes in the expression of *Cd8a, Cd68*, and *Nkp46*, immune cell markers for cytotoxic T-cells, macrophages, and natural killer cells, respectively. *Cd8a* showed a 6-fold increase peaking after 2 cycles then decreased with further CPA treatment, perhaps reflecting CPA-induced immune cell cytotoxicity. Similarly, *Cd68* increased 3-fold after 4 cycles then declined, while *Nkp46* showed no significant changes in expression at any time point (Fig. 4C). The immune cell effector markers *Ifng, Prf1* and *Gzmb* also showed peak induction after 2 CPA cycles then decreased with further treatment.

**Fig. 4.**
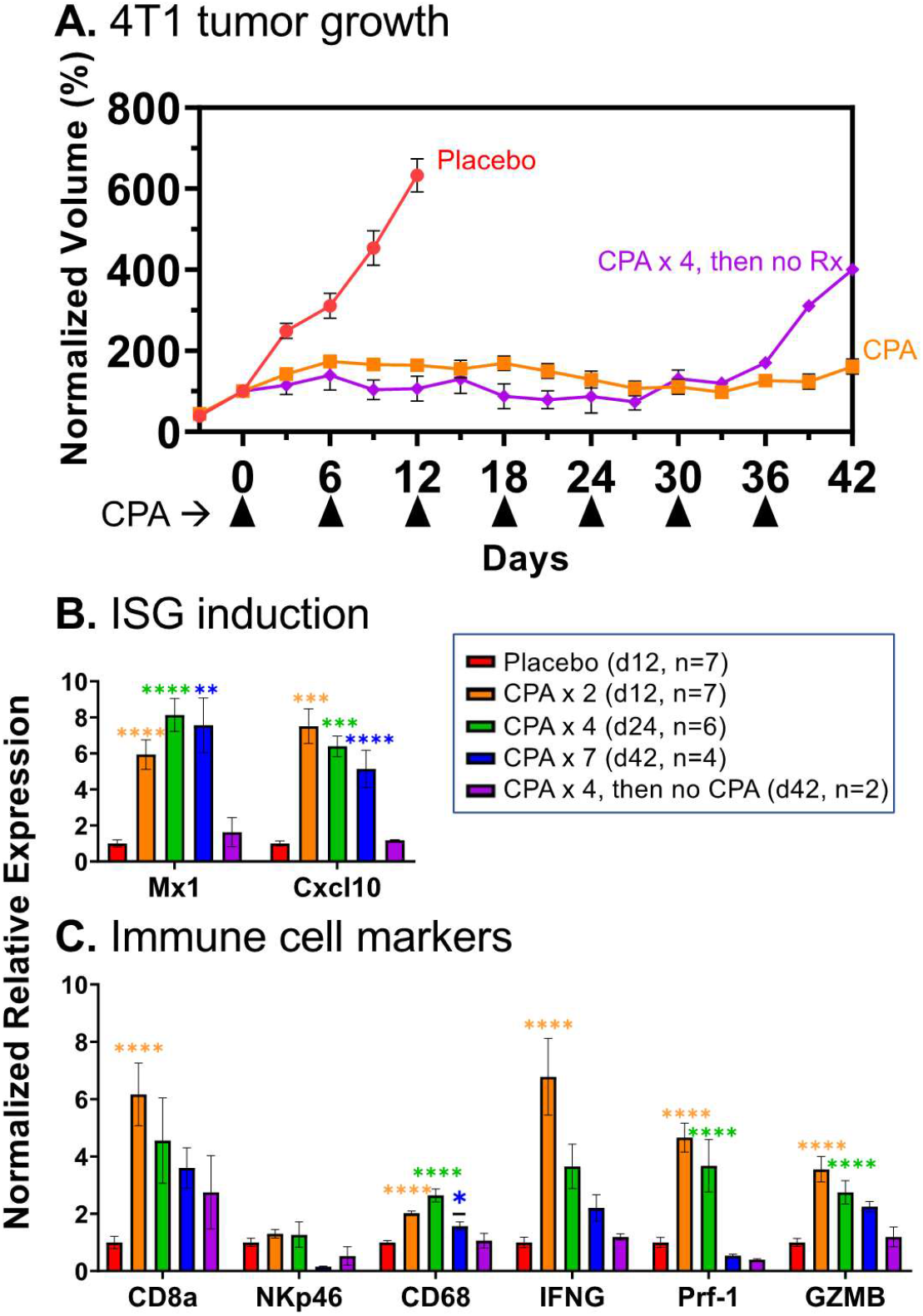
4T1 tumor growth and gene expression changes induced by CPA treatment. **A**. 4T1 cells were implanted orthotopically in 6-week-old female BALB/c mice, then treated with 130 mg/kg CPA or PBS (placebo) on a 6-day metronomic schedule once mean tumor volumes reached 100-150 mm^3^. Shown are group tumor volumes (mean +/-SEM) normalized to the volume on the first day of CPA treatment (day 0). Mice were euthanized and tumors excised for qPCR analysis of tumor RNA (**B, C**) on treatment days 12, 24 and 42, with n tumors/group, as indicated (box). Thus, the curve marked CPA represents 17 tumors through day 12, then 10 tumors through day 24, and then 4 tumors through day 42 (7 CPA treatment cycles). Data indicate exponential growth of placebo group vs growth stasis through 7 CPA treatment cycles. Tumors began to regrow by day 36 when CPA was halted after 4 treatment cycles. **B, C**. qPCR analysis of ISGs and immune cell marker genes in tumor cell RNA. CPA induced tumor ISG expression after 2, 4 and 7 treatment cycles, but the induction was reversed when treatment was halted after 4 cycles. CPA induction of cytotoxic effector expression and immune cell infiltration were reduced or lost with prolonged treatment. Significance (1-way ANOVA): *, p < 0.05; **, p < 0.01; ***, p < 0.001; ****, p < 0.0001.

These findings were validated by RNA-seq analysis of total tumor RNA across the time course, which identified hundreds of treatment-induced gene responses. Innate immunity, immune response and cellular response to interferon-beta represent the top Functional Annotation Clusters of up regulated genes after 2, 4 and 7 CPA treatment cycles (Table S4). Notably, these immune response gene clusters were not found in the regrowing tumors when CPA treatment was halted after 4 cycles (Table S4). Thus, in 4T1 tumors, metronomic CPA activated a transient immune response associated with tumor growth stasis.

### Metronomic CPA induces robust E0771 tumor regression and immune cell recruitment

Major regression of E0771 tumors was seen within 2 CPA treatment cycles (Fig. 5A), in contrast to the growth stasis response of 4T1 tumors. ISG induction was comparatively weak; it was first seen on day 2, peaked on day 3, then declined and was undetectable by day 12 (Fig. 5B). Importantly, ISG induction was followed by strong immune cell recruitment by days 6 and 12 (Fig. 5C). T-regs (*Foxp3*) initially decreased at day 2 before returning to basal levels by day 6 (Fig. 5C).

**Fig. 5.**
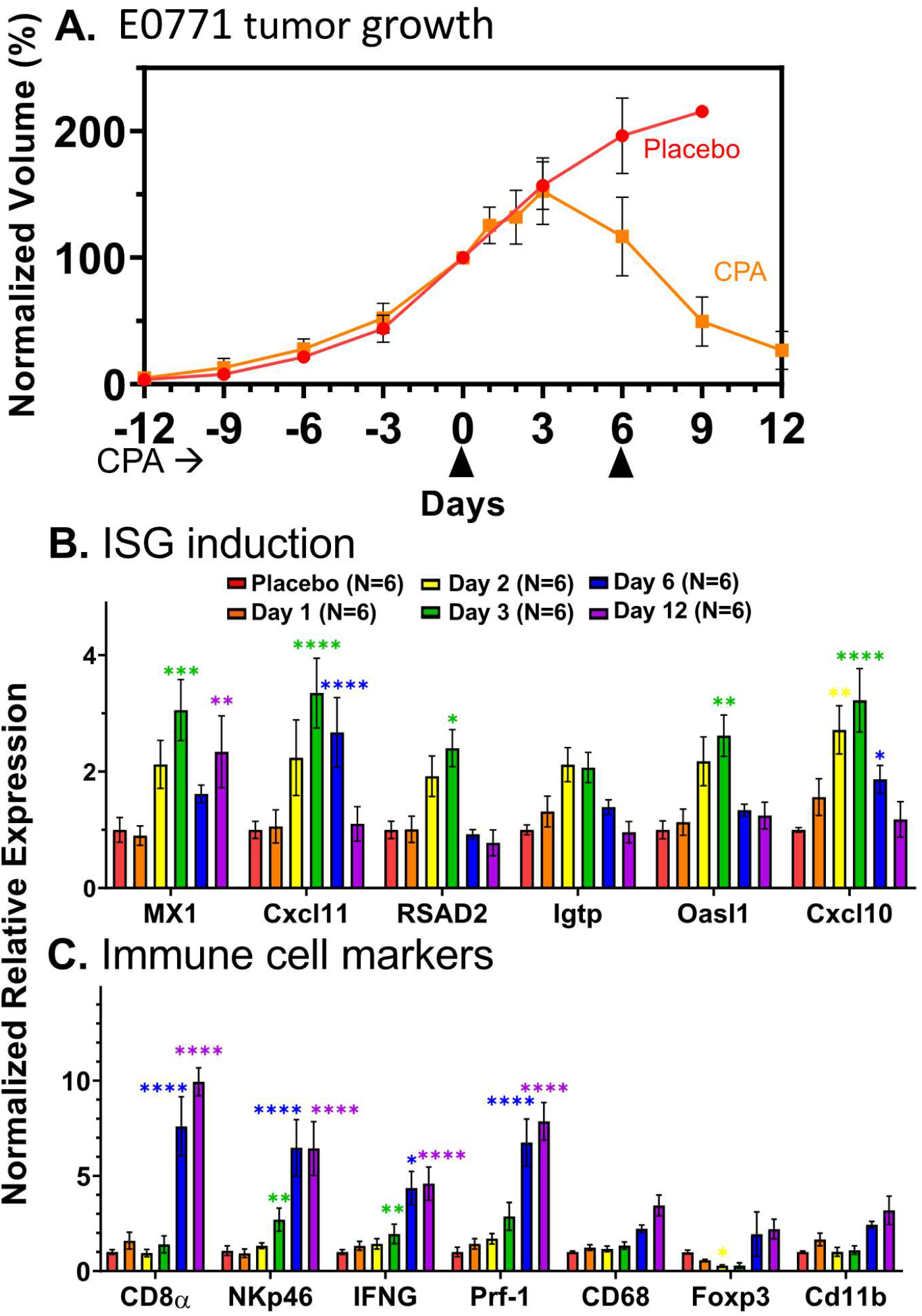
Tumor growth and gene expression changes in CPA-treated E0771 tumors. **A**. Impact of metronomic CPA treatment (110 mg/kg per injection; arrow heads along x-axis) on E0771 tumor growth. Data shown are mean +/-SEM tumor volumes for n=6 tumors per group, normalized to 100 percent of the Day 0 volume. **B, C**. qPCR analysis as in Fig. 4. The weak induction of ISGs was highest three days after the first CPA dose, at which time immune cell infiltration and cytotoxic effectors were first increased and then maintained through day 12. Increases in macrophages and dendritic cells (*Cd11b*) were significant by t-test but not by ANOVA. Significance vs placebo group (2-way ANOVA): *, p < 0.05; **, p < 0.01; ***, p < 0.001; ****, p < 0.0001.

E0771 tumor RNA-seq analysis revealed an interesting pattern: there were relatively few gene expression changes during the first 3 days after CPA treatment, followed by large numbers of treatment-responsive genes on days 6 and 12 (i.e., after 1 and 2 treatment cycles) (Fig. 6A). Genes were grouped by whether their response to treatment was early (days 1-3) or late (days 6, 12), and whether the response was sustained through day 12 or was not (i.e., transient) (Fig. 6B, Table S5A, S5B). Top Functional Annotation Clusters included innate immunity for both Early-Transient and Late up regulated genes, whereas inflammatory response was a top cluster term for Early-Sustained induced genes (Table S5C-S5F). Comparison to the set of 188 type-I IFN response genes identified in cultured E0771 cells (Fig. 3A; common response to 4HC and IFNB) revealed a striking, 61-fold enrichment in the set of 73 Early-Transient (induced) genes, 52 (71%) of which were in the 188 gene set (p < E-05 vs background set of all expressed genes; Fisher’s exact test). The type-I IFN response genes showed no enrichment in the Early-Sustained induced gene set (0 of 56 genes) and marginal enrichment in the Late induced gene set (35 out of 1,666 genes; 1.38-fold enrichment, p = 0.09) (Fig. 6B). Thus, CPA induces an early type-I IFN response that is not sustained through day 12, by which time there is major immune cell infiltration (Fig. 5C) and a 14-fold increase in the overall number of differentially expressed genes (Fig. 6A). Early-Sustained down regulated genes were enriched for sterol and lipid metabolism, while the large set of Late down genes was enriched for cell cycle and transcriptional regulation terms (Fig. S5G, S5H).

**Fig. 6.**
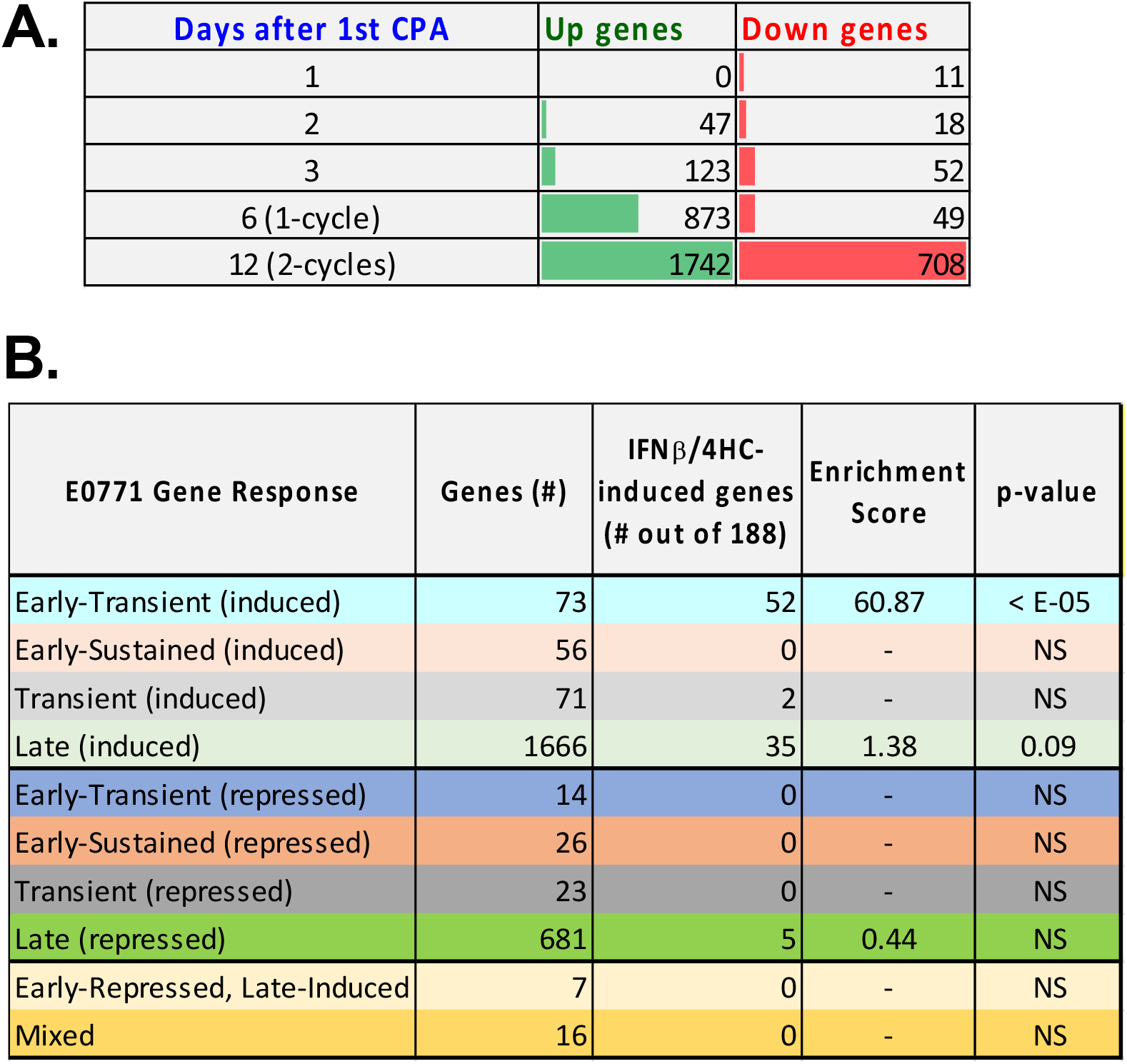
E0771 tumor RNA-seq. **A**. Number of genes showing significant up regulation or down regulation at each of 5 time points of metronomic CPA treatment. A total of 2,633 genes met the thresholds for a significant response (Fold-change > 2 at edgeR-adjusted p-value < 0.05) at one or more time points (Table S5A). **B**. The set of 2,633 responsive genes was classified based on the time course of response, as detailed in Table S5B. Each set was analyzed for overlap with the set of 188 genes that were up regulated by both 4HC and IFNB in cultured E0771 cells (Fig. 3B), and enrichment scores with significance by Fisher exact text calculated compared to a background set of all genes expressed at FPKM > 1 as shown in Table S5B.

Finally, ALAS2, a heme biosynthetic enzyme, and six hemoglobin genes (most notably four HBB genes) were strongly down regulated by CPA on days 1-3, but then strongly up regulated after 2 treatment cycles (Fig. 6B, Table S5I). Hemoglobin-beta (HBB) contributes to breast cancer neoangiogenesis and metastasis by a tumor cell protective anti-oxidant mechanism (37,38), but also becomes a dominant self-antigen target of CD8-T cells in tumor pericytes following IL12 immunotherapy (39).

### E0771 tumor regression requires CD8+ T-cells

Next, we used an immune-depletion strategy to ascertain the role of CD8-T cells in metronomic CPA-induced E0771 tumor regression. To minimize the direct effects of CPA cytotoxicity and maximize possible immune system contributions, we decreased the CPA dose from 130 mg/kg to 110 mg/kg, which was effective at inducing tumor regression, ISG induction, and immune cell recruitment (Fig. S4). Anti-mouse CD8α antibody, or control IgG, was administered to E0771 tumor-bearing mice beginning 5 days before the first CPA treatment on day 0.

FACS analysis confirmed the depletion of circulating CD8 T-cells by day 1, which persisted for at least 5 weeks after the last antibody injection on day 15 (Fig. S5, Fig. S6). Moreover, in contrast to the near-complete tumor regression achieved in the CPA + control IgG group, an extended period of tumor growth stasis followed by robust tumor regrowth was evident in mice receiving CPA + anti-CD8α antibody (Fig. 7A). We conclude that CD8 T-cells are essential for CPA-induced tumor regression, and in their absence, E0771 tumors escape the cytotoxic effects of CPA treatment.

**Fig. 7.**
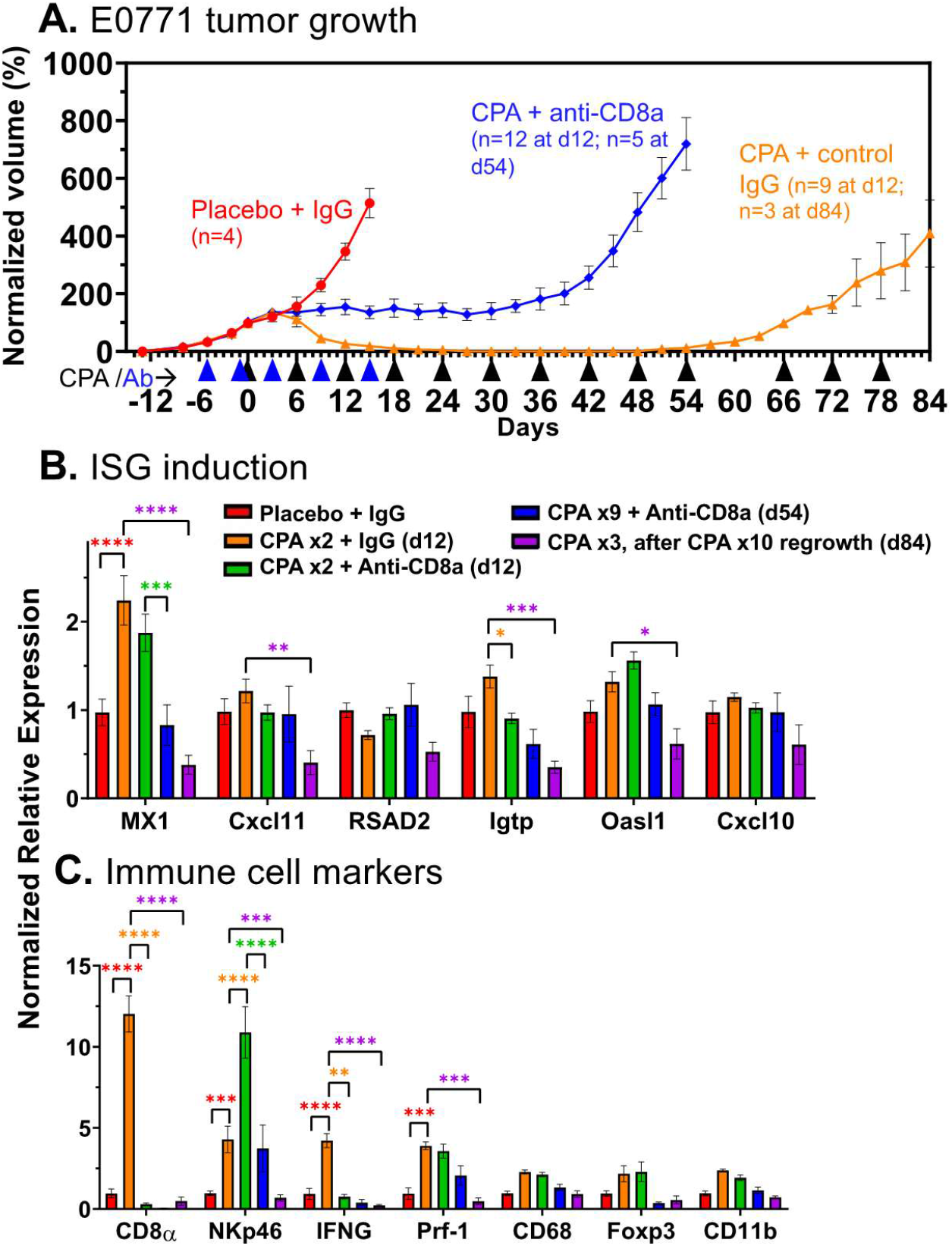
Impact of CD8a immunodepletion on CPA-induced E0771 tumor regression and immune cell recruitment. **A**. E0771 tumors were treated with CPA every 6 days at 110 mg/kg (black arrow heads along x-axis), alone or in combination with anti-CD8a or control IgG (blue arrow heads). Tumor volumes were normalized to the percent of Day 0 volume (=100). By day 12, the placebo + control IgG, CPA + control IgG, and CPA + Anti-CD8a groups showed 3 distinct growth patterns: exponential growth, tumor stasis and tumor regression, respectively. Tumors resumed growth by day 42. CPA-regressed tumors eventually regrew and became resistant to CPA treatment. Data shown are mean +/-SEM volumes for n=4 tumors for the placebo + IgG group, n=5 for CPA + IgG (d12), n=3 for CPA + IgG (d84), n=7 for CPA + anti-CD8a (d12) and n=5 for CPA + Anti-CD8a (d54). **B**. ISG induction was weak in all groups, but the regrowing CPA-resistant tumors showed decreased ISG expression. **C**. Anti-CD8a antibody prevented CPA-induced CD8 T-cell infiltration and IFNG production, but NK cell infiltration increased. Significance (2-way ANOVA): *, p < 0.05; **, p < 0.01; ***, p < 0.001; ****, p < 0.0001.

CPA induction of the ISG *Mx1* was unaffected by anti-CD8α antibody, as was expected given the expectation that ISG gene induction occurs upstream of immune cell infiltration. Other ISGs, whose induction by CPA in E0771 tumors was transient (seen on days 2, 3 and 6, but not day 12; Fig. 5B), were not induced in the day 12 tumor samples (Fig. 7B). Importantly, anti-CD8α antibody abolished the increase in tumor-infiltrating CD8 T-cells (*Cd8a*), as well as the increase in *Ifng*, which is produced by tumor-infiltrating CD8 T-cells and could be an important contributor to tumor regression. Surprisingly, the induced expression of the NK cell marker *Nkp46* was further augmented by anti-CD8α, while *Prf1*, which is produced by both CD8 T-cells and NK cells, showed no net change in expression (Fig. 7C).

Regrowth of the regressed CPA + control IgG treated tumors became apparent once CPA treatment was discontinued on day 60, after which the tumors became resistant to further CPA treatment (Fig. 7A). The expression of *Mx1* decreased below basal levels in the regrowing tumors (Fig. 7B), as did that of *Cd8a, Nkp46* and the cytotoxic effectors *Ifng* and *Prf1* (Fig. 7C), which may contribute to the emerging resistance to CPA. Together, these findings provide strong support for the conclusion that metronomic CPA-induced recruitment of CD8 T-cells is essential for E0771 tumor regression.

### Role of type-I IFN signaling in CPA-induced immune cell recruitment and tumor regression

We used an inhibitory IFNAR-1 antibody to determine the functional role of type-I IFN signaling and the impact of the transient, downstream induction of ISGs on CPA-induced immune cell recruitment and tumor regression. Mice bearing E0771 tumors were given anti-mouse IFNAR-1, or control IgG, beginning 1 day prior to the first CPA treatment on day 0 (Fig. 8A). Remarkably, the major tumor regression seen by day 12 in the CPA + control IgG mouse group was fully blocked in all 16 mice given CPA + anti-IFNAR-1 antibody. Furthermore, robust tumor growth persisted in 5 of the 8 mice that we continued to monitor after anti-IFNAR1 treatment was halted on day 12, and only moderate regression was observed in the 3 other mice, despite ongoing CPA treatment. This result contrasts to the growth static response to CPA seen in CD8 T-cell-depleted tumors (Fig. 7) and indicates that direct CPA tumor cell cytotoxicity has limited impact on tumor growth in the absence of IFNAR1 signaling.

**Fig. 8.**
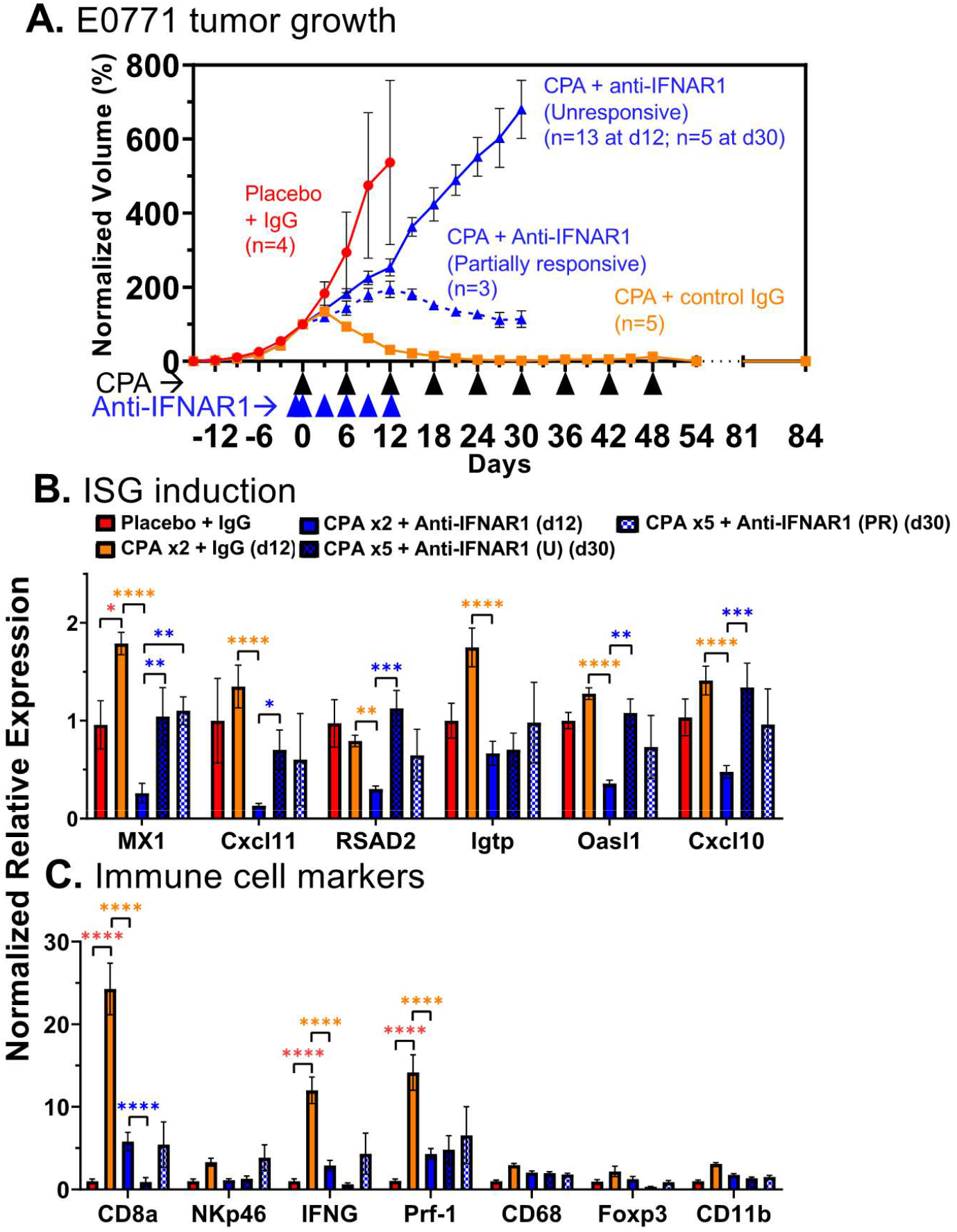
Type-I interferon signaling is required for E0771 tumor regression. **A.**E0771 tumors were treated with metronomic CPA, as in Fig. 6, alone or in combination with anti-IFNAR1 to block type-I interferon signaling. Data shown are mean +/-SEM volumes for n=4 tumors for the placebo + IgG group, n=5 for CPA + control IgG, and n=16 (through day 12) for CPA + anti-IFNAR1 (decreasing to n=5 from day 12-30, due to 8 tumors excised for analysis on day 12). Tumors in the CPA + anti-CD8a group showed two distinct growth patterns: Unresponsive, with strong continued growth through day 30; and Partially responsive, as indicated by growth stasis or moderate regression, as marked. Tumor volumes were normalized to the percent of Day 0 volume (=100). Significance (2-way ANOVA): *, p < 0.05; **, p < 0.01; ***, p < 0.001; ****, p < 0.0001. **B, C**. Tumor ISGs were significantly depleted by anti-CD8a antibody, which also suppressed CPA-induced CD8 T-cell infiltration and IFNG and Prf-1 production. Of note, in this cohort of mice, the CPA + control IgG tumors were eradicated and by day 84 did not show the tumor regrowth seen in the mice shown in Fig. 7A.

Gene expression analysis showed that anti-IFNAR1 antibody reduced ISG expression below control tumor levels by day 12, indicating the antibody is highly effective in blocking tumor IFN signaling (Fig. 8B). ISG expression returned to basal levels by day 30, i.e., 18 days after antibody treatment was halted on day 12. Anti-IFNAR1 also reduced tumor infiltration of all tested immune cells by day 12 (Fig. 8C).

The reduction of tumor infiltrating CD8 T-cells was further supported by FACS analysis of tumor tissue (Fig. S7A) and occurred without any changes in circulating CD8 T-cells (Fig. S7B). Thus, depletion of tumor infiltrating CD8 T-cells, and likely other infiltrating immune cells, is a consequence of the inhibition of tumor type-I IFN signaling and not a systemic effect. Day 12 CD8a T cell marker levels were restored by day 30 in the three anti-IFNAR1-treated tumors that were partially responsive to CPA, but not in the five CPA-unresponsive tumors (Fig. 8C, PR vs. U groups), which could help explain their differences in CPA responsiveness.

## Discussion

Effective treatment of TNBC continues to be challenging, with limited therapeutic options and high rates of disease recurrence (40). Cytotoxic chemotherapies, including CPA, remain the primary systemic treatment modality for TNBC despite highly variable treatment responses and frequent development of chemo-resistance (41). Here, we explored the role of innate immunity in relation to cytotoxic treatment response and resistance in TNBC. We used two orthotopic mouse models of TNBC, 4T1 and E0771, to investigate the chemo-immunogenic activity of CPA when delivered on a metronomic, medium-dose intermittent schedule (22). 4HC, a chemically activated CPA derivative that spontaneously decomposes to yield the same active metabolite as CPA, induced the expression of hundreds of ISGs in both TNBC cell models in a manner similar to doxorubicin, an established immune-stimulatory chemotherapeutic agent (42). These tumor cell-centric ISG responses to activated CPA were at least in part dependent on signaling by the type-I IFN receptor, IFNAR-1, implicating tumor cell production of type-I IFNs in these drug-induced ISG responses. Many of the ISG responses seen in TNBC cell culture were recapitulated *in vivo* in MEDIC CPA-treated TNBC tumors implanted in syngeneic mice. Notably, CPA-treated 4T1 tumors showed robust type-I IFN signaling and tumor immune cell infiltration, leading to an overall tumor growth static response. In contrast, E0771 tumors exhibited a somewhat weaker IFN response, but this was followed by robust immune infiltration and extensive tumor regression, both of which were absolutely dependent on type-I IFN signaling by IFNAR-1. Thus, a robust IFN-mediated immune response may be essential for the efficacy of metronomic CPA in TNBC. Furthermore, our findings raise the possibility that treatment resistance to CPA, and perhaps other chemo-immunogenic cytotoxic agents, may stem from silencing of the IFN pathway.

ISG induction was an early response to drug treatment in both TNBC models, both in cell culture and following CPA treatment of implanted tumors *in vivo*. The ISG responses seen *in vivo* were transient (Fig. 6B) and were followed by strong increases in both innate and adaptive infiltrating immune cells, including NK cells and CD8a T-cells, after 6-12 days (i.e., 1-2 CPA treatment cycles). Type-I IFNs and the ISGs they induce are known to stimulate T-cells, NK cells, macrophages and dendritic cells, and other immune cells (18). ISG induction may thus be a useful marker for immunogenic potential *in vivo*. Of note, the ISG responses seen in our TNBC models were not apparent until 48 h after drug treatment, even though ISG gene induction *per se* is a rapid process, as was seen when TNBC cells were treated with IFNβ directly (Fig. S3). The delay in ISG induction seen in CPA-treated TNBC cells and tumors likely reflects time required for CPA to effect tumor cell damage and the associated production of immunostimulatory damage-associated molecular pattern molecules. These may include double stranded RNAs and nucleic acid agonists of STING, which can activate cytosolic sensors and induce type-I IFN production through established mechanisms (21).

Using RNA-seq, we validated the transient nature of CPA-induced ISG responses on a global scale. We identified a set of 188 ISGs that responded in common to IFNβ and to 4HC treatment in cultured E0771 cells, as well as 380 commonly responding ISGs in 4T1 cells. Strikingly, 52 of the 188 E0771 ISGs showed an early, transient response in CPA-treated E0771 tumors, where they comprised 71% of the Early-Transient response gene set, representing a 61-fold enrichment compared to a background set of all expressed genes (Fig. 6). These 52 genes comprise a robust set of CPA-responsive E0771 ISGs and could serve as useful markers for early chemo-immunogenic responses to CPA treatment *in vivo*.

The ability of anti-IFNAR1 antibody to almost completely abolish CPA-induced E0771 tumor regression establishes that type-I IFN signaling is essential for the anti-tumor actions of metronomic CPA in this TNBC model. This, in turn, leads us to the unexpected conclusion that the intrinsic tumor cell cytotoxicity of CPA does not translate into a major therapeutic response in the absence of type-I IFN signaling.

These findings support a model whereby CPA-induced tumor cell damage induces the major anti-tumor effects of CPA on E0771 tumors indirectly, via its ability to activate tumor cell autonomous type-I IFN signaling linked to an immunogenic cell death mechanism. Of note, we observed tumor growth stasis was when circulating and tumor cell infiltrating CD8 T-cells were immuno-depleted, i.e., the block in CPA anti-tumor activity was less complete than with anti-IFNAR1 antibody. This tumor growth stasis response is likely mediated by other tumor infiltrating immune cells, e.g., NK cells, whose CPA-induced levels were further increased by CD8a T-cell depletion (Fig. 7C). The ability of NK cells to contribute to the anti-tumor effects of metronomic CPA is supported by earlier findings in glioma models (24,43).

The two TNBC models studied here, 4T1 and E0771, exhibited notable differences in their ISG, immune cell, and therapeutic responses to MEDIC CPA treatment *in vivo*. 4T1 tumors were characterized by a stronger and longer lasting ISG induction, but this did not translate into a greater anti-tumor response.

This is evidenced by the growth stasis observed in 4T1 tumors versus the major regression seen in E0771 tumors. Combination chemo-immuno therapies designed to stimulate immunogenic cell death and activate a more robust anti-tumor response (44) may be required for more effective treatment of 4T1 tumors. Differences in mouse strain, tumor cell proliferation and angiogenesis, and mutational burden, which is much higher in E0771 than 4T1 tumors (45), could contribute to the differential responsiveness of these two TNBC models to CPA treatment. In addition, immunosuppressive regulatory T-cells (Foxp3^+^ CD4^+^) increase with time in both TNBC models, but only E0771 tumors display an early growth period when a majority of tumor-associated CD4^+^ T-cells are immunostimulatory (29), resulting in a more favorable environment for CPA responses. Further study of the mechanisms underlying metronomic CPA-induced E0771 tumor regression and the comparative resistance of 4T1 tumors may help identify useful biomarkers for tumor responsiveness and could lead to the discovery of new molecular targets for increasing effectiveness of chemotherapy in poorly responsive TNBC. Recent clinical trials have used CPA in combination with other drugs to treat TNBC with varying degrees of success (46-49), and there may be opportunities for further improvements based on metronomic dose and schedule optimization (22).

Finally, the treatment models developed here may provide an important means to develop clinically translatable markers of chemo-immunogenic treatment response and resistance. The gene signatures identified by our RNA-seq analysis may be useful in pre-treatment biopsies to identify patients more likely to elicit an IFN-mediated treatment response. There is also substantial interest in identifying non-invasive imaging markers of chemotherapy treatment response (50). Therapy-induced responses, including apoptosis, proliferation, and overall treatment response, can be monitored in real time in preclinical oncology models by label-free optical imaging techniques, such as spatial frequency domain imaging (51,52).The preclinical therapy models described here provide a means to discover novel imaging markers that discriminate chemo-immunogenic sensitive and resistance tumors (53). As similar imaging markers can be tracked in patients with clinical imaging modalities such as PET and optical diffuse optical spectroscopy, it may be possible to rapidly identify both treatment response and resistance and adjust the treatment regimen accordingly (54,55).

## Author contributions

Initial 4T1 cell culture work and 4T1 tumor model studies were performed by KAD and CV. All of the other experimental work, including studies of cultured E0771 cells and tumors and RNA-seq analysis was carried out by CV. Data analysis and preparation of figures were carried out by CV and DJW. The manuscript was initially drafted by CV with input from DJW and DR, then edited and finalized by DJW. All authors contributed to experimental design and reviewed and approved the final manuscript. The project was conceived and supervised by DJW.

## Supplemental figures

**Fig. S1.**
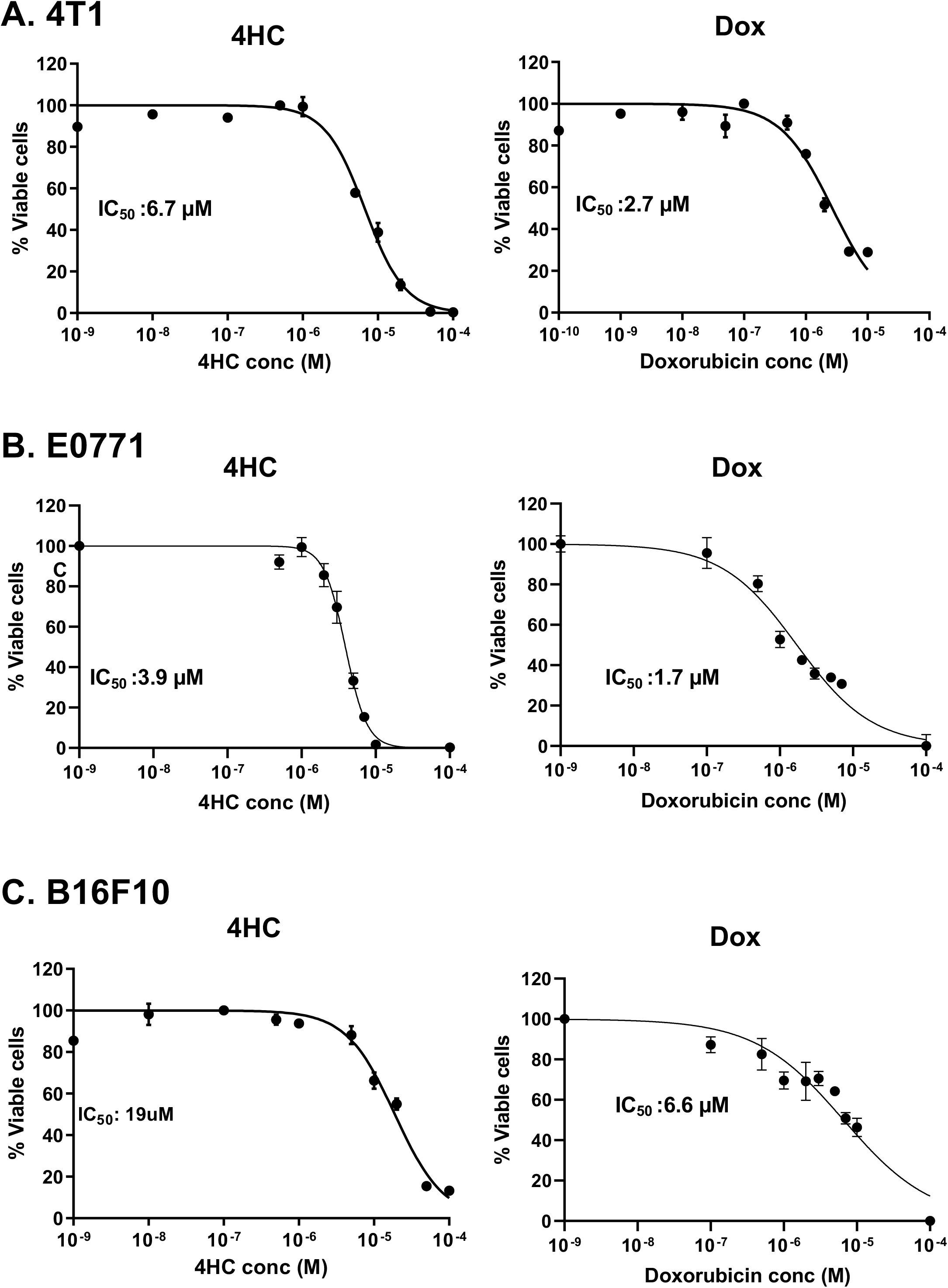
Dose-dependence of drug sensitivity of cultured 4T1, E0771 and B16F10 cells. Shown are viability assays determined in MTS assays over multi-log_10_ range of 4HC (left) and doxorubicin (right) for 4T1 (**A**), E0771 (**B**) and B16F10 cells (**C**). Data points: mean +/-SD values for n = 3 wells of a 96-well plate. IC50 values were determined using log (inhibitor) vs normalized response function in GraphPad Prism.

**Fig. S2.**
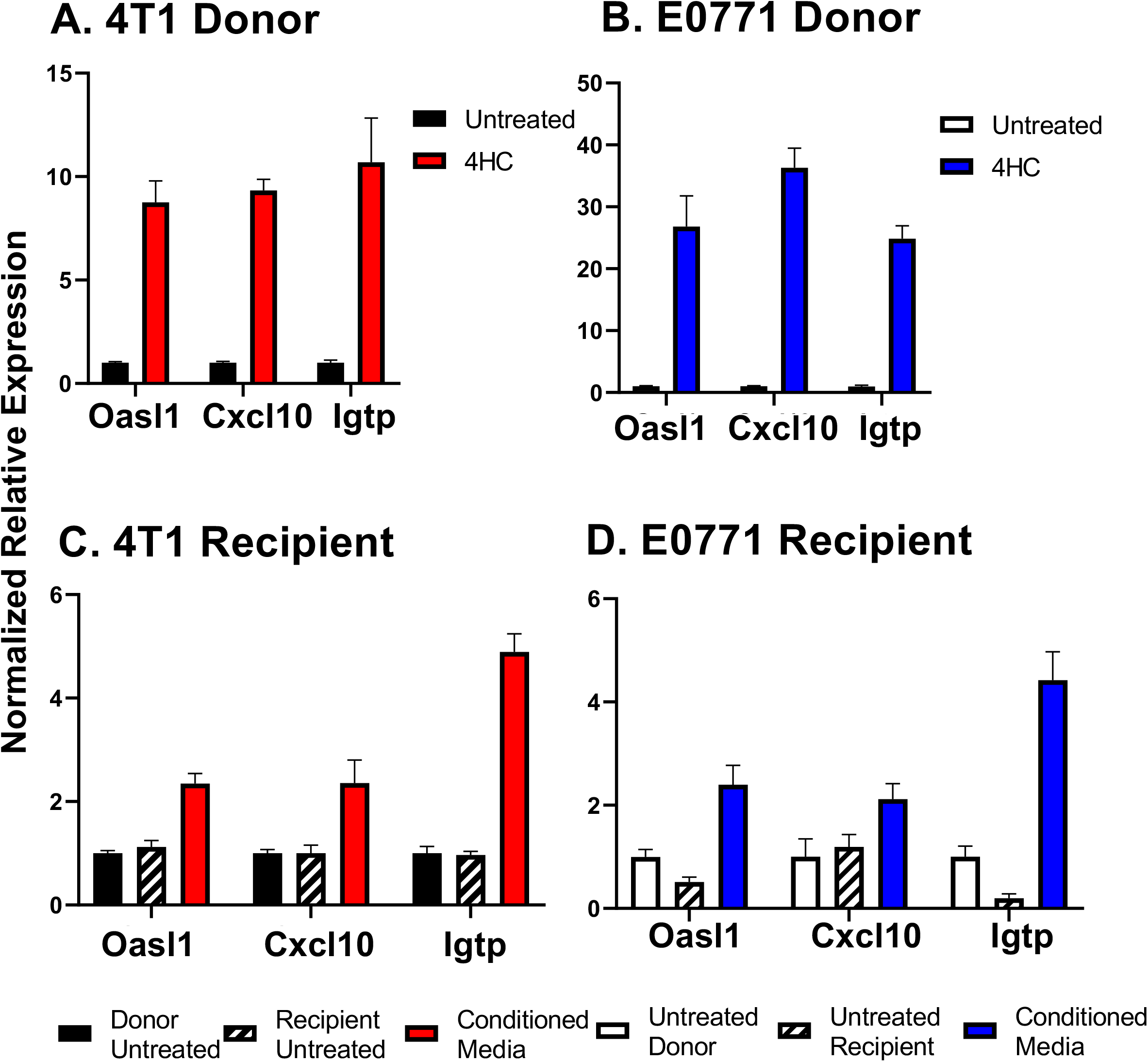
ISG induction by 4HC-conditioned culture medium. **A, B**. 4T1 and E0771 cells were treated with 4HC for 72-h under the same conditions as Fig. 1. Data show ISGs were strongly induced in both cell models, as determined by qPCR analysis. **C, D**. Induction of ISGs in drug-naïve 4T1 and E0771 recipient cells treated for 4-h with 4HC-conditioned cell culture supernatant from the corresponding drug-treated donor cells (as in A, B), followed by a PBS wash and 2 h incubation in fresh culture medium. Gene expression was analyzed by qPCR. In both cell lines, recipient cells showed weaker ISG induction than in cells directly exposed to 4HC. Data points: mean +/-SD values for n = 2-3 replicates representative of two independent experiments.

**Fig. S3.**
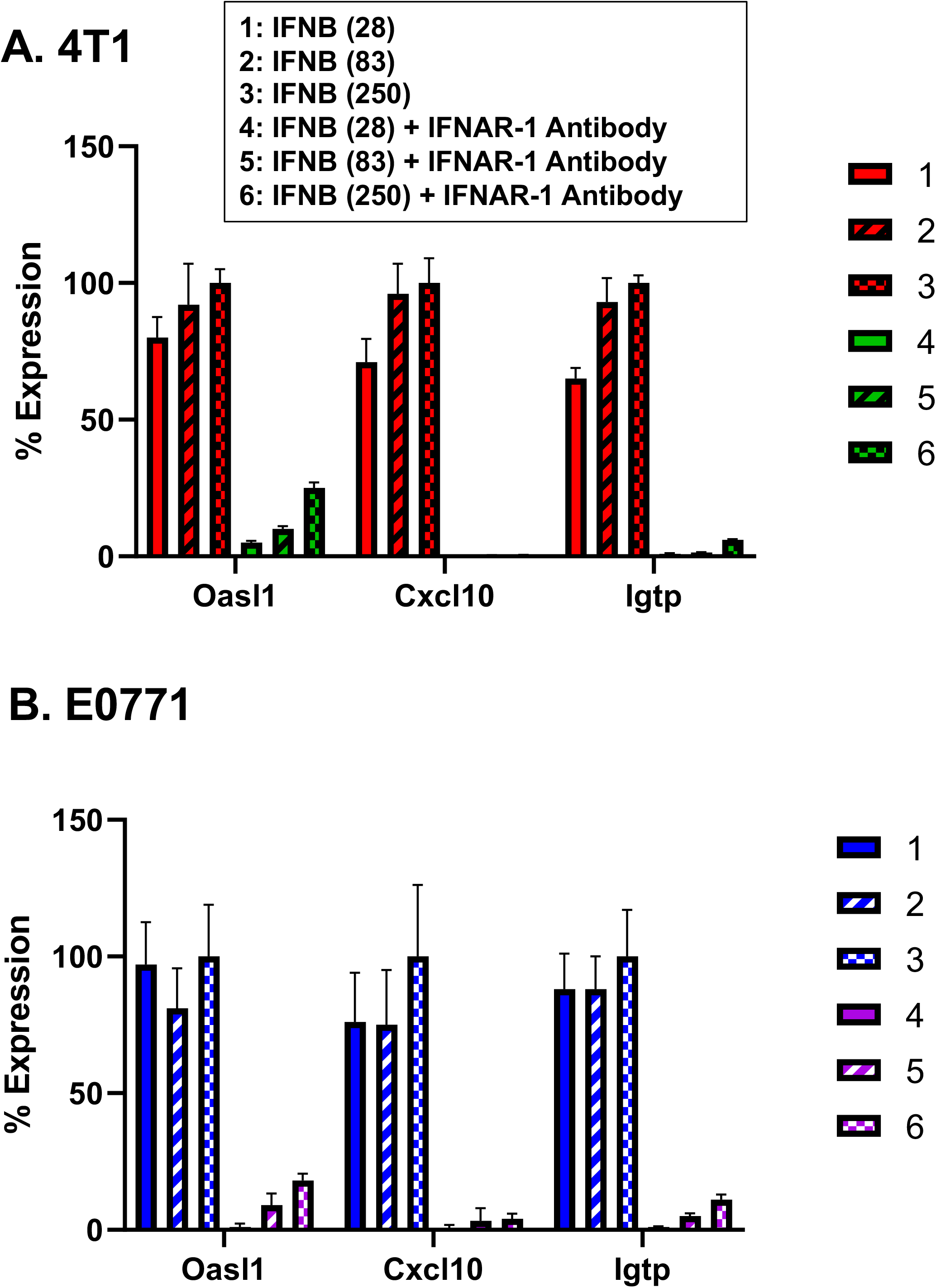
Verification of anti-IFNAR-1 antibody inhibitory activity. 4T1 cells (**A**) and E0771 cells (**B**) were treated with 28, 83 or 250 U/mL of mouse recombinant IFNβ for 4-h, with or without 10 µg/mL IFNAR-1 antibody and harvested 2 h later. IFNβ induced similar ISG responses in both cell lines at all concentrations. Anti-IFNAR1 antibody blocked ISG induction > 90%, except at the highest concentration IFNβ for the ISG *Oasl1*, where the antibody concentration may not have been sufficient to effect complete inhibition. Data presented as mean +/-SD with n=3 replicates. Percent expression was calculated using the formula ((x ± SD) - (z ± SD)) / ((y ± SD) - (z ± SD)), where × = antibody + IFNβ treatment gene expression, z = untreated gene expression and y = IFNβ treatment gene expression.

**Fig. S4.**
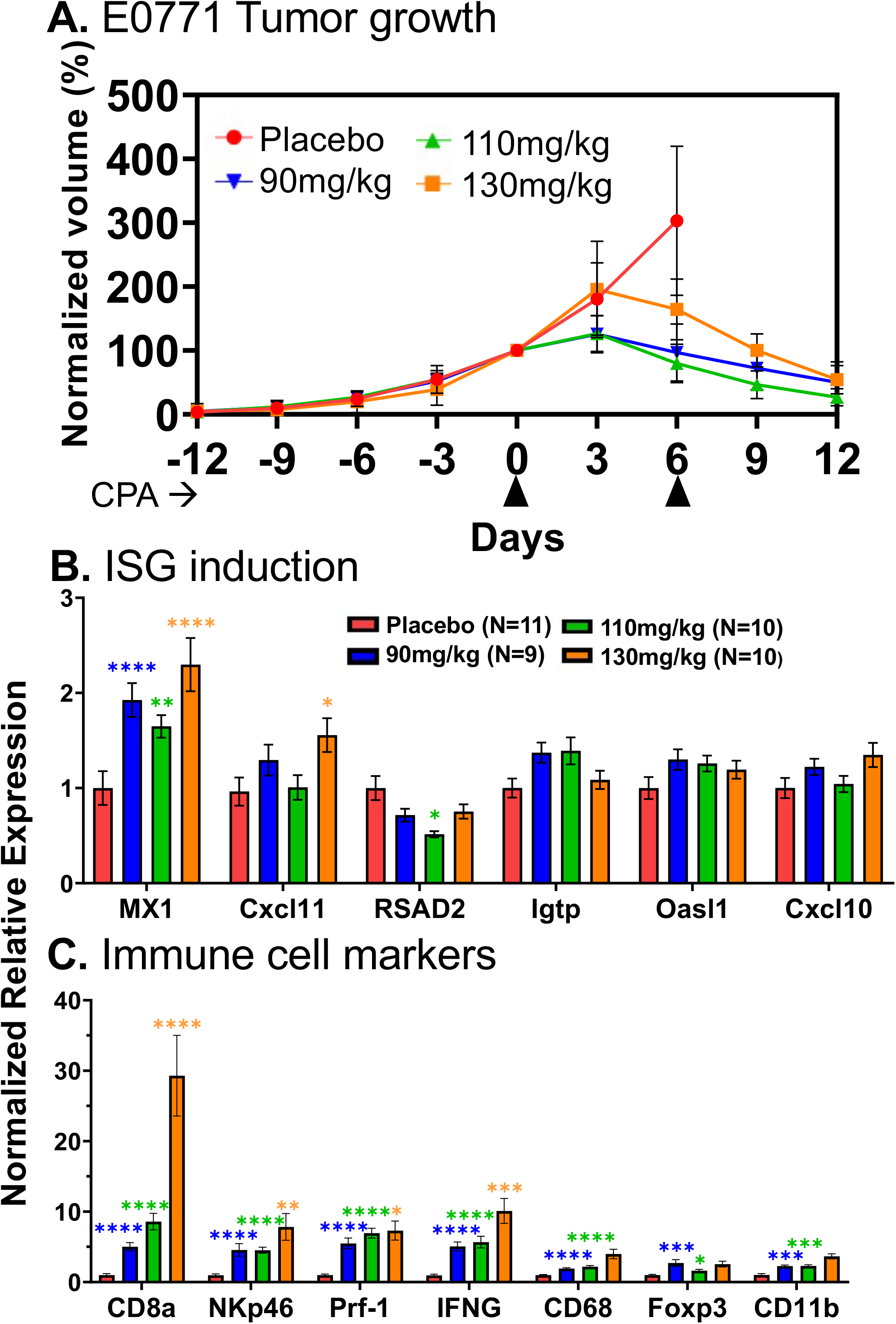
E0771 tumor growth curves and qPCR analysis of metronomic-CPA *in vivo* dose response data. **A**. E0771 tumors implanted in mice were treated with 90, 110 or 130 mg/kg CPA every 6 days. All three CPA dosages induced extensive tumor regression by day 12. Data shown are mean +/-SEM values for n = 11 tumors for the placebo group, n = 9 tumors for the 90 mg/kg CPA group, n = 10 tumors for the 110 mg/kg CPA group, and n = 10 tumors for the 130 mg/kg CPA group. Tumor volumes were normalized to 100 percent of the volume on Day 0 (first day of CPA treatment).**B**. ISG induction was < 2-fold at all CPA doses. **C**. Immune cell marker genes showed very similar fold-change values at each CPA dose, except for CD8a, which showed dose-dependent induction. Significance was determined by 2-way ANOVA: *, p < 0.05; **, p < 0.01; ***, p < 0.001; ****, p < 0.0001.

**Fig. S5.**
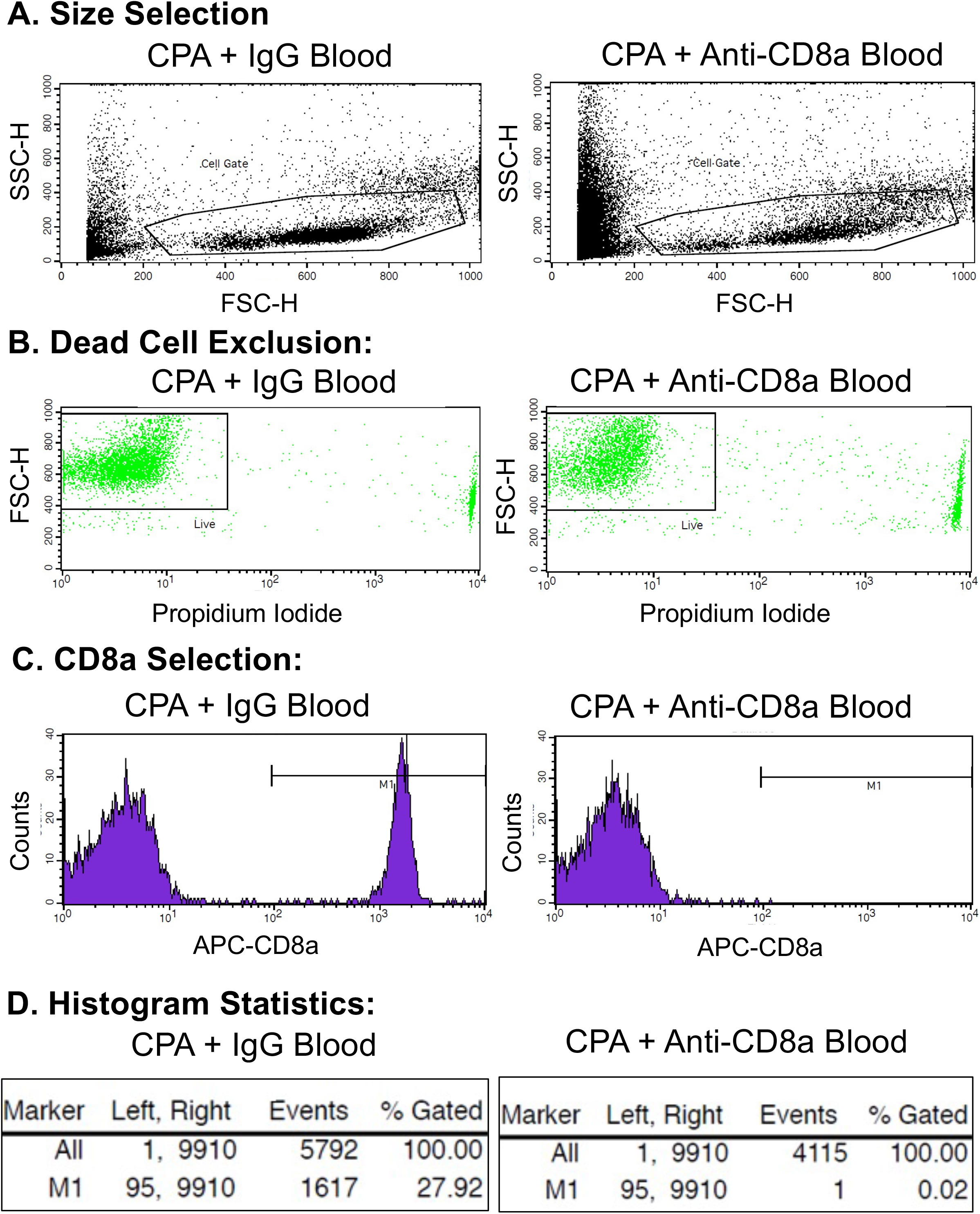
Representative FACS analysis of blood from CPA-treated mice, with and without anti-CD8a antibody treatment. Data representative of 2 individual mice. **A**. 20 uL of mouse tail vein blood was prepared for FACS analysis of circulating CD8 T-cells. Events were selected based on general size parameters of forward-scatter (FSC-H) and side-scatter (SSC-H) to exclude overly large and small events. **B**. Live cells were selected by excluding events with propidium iodide signal. **C**. CD8 T-cells were selected by excluding events that lacked the APC signal from the APC-labeled anti-CD8a antibody used in sample preparation. **D**. CD8 T-cell percentages were calculated by dividing the CD8+ events by the total number of live events. Mouse blood from the CPA + anti-CD8a group was devoid of CD8 T-cells, in contrast to blood from the CPA + IgG control group.

**Fig. S6.**
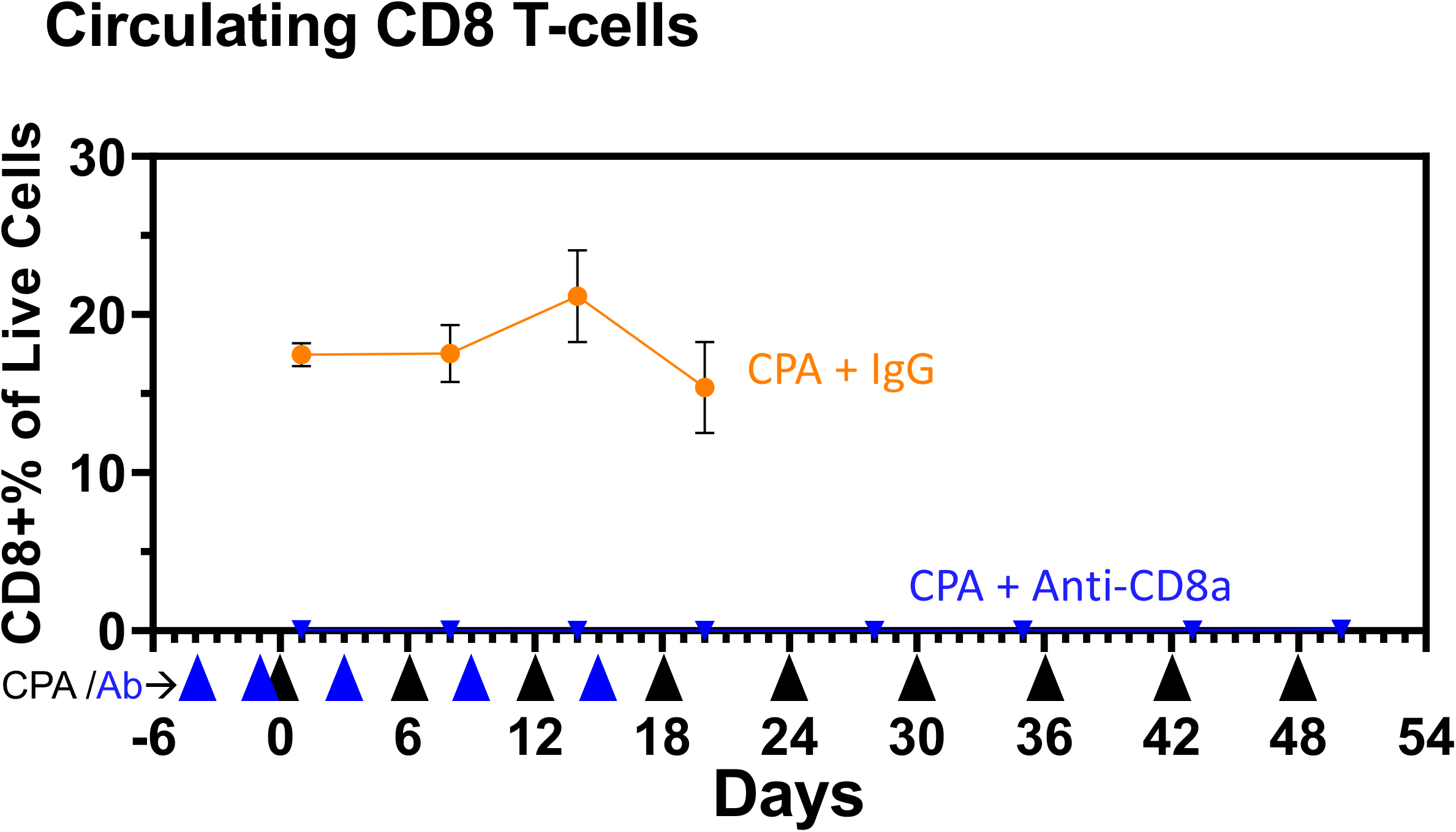
Circulating CD8 T-cells for CPA-treated mice with and without anti-CD8a antibody. FACS analysis of blood from the mice shown in Fig. 7 that were given CPA + anti-CD8a antibody showed complete depletion of circulating CD8 T-cells after 2 antibody doses. The depletion was maintained for many weeks after antibody treatment (blue arrow heads below x-axis) was halted. CD8 T-cells levels were not detected in the CPA + anti-CD8a group at any of the 8 time points analyzed (small triangles superimposed on the x-axis, at time points from day 1 through day 50). Data shown are group mean values +/-SEM for n = 4 mice in each group.

**Fig. S7.**
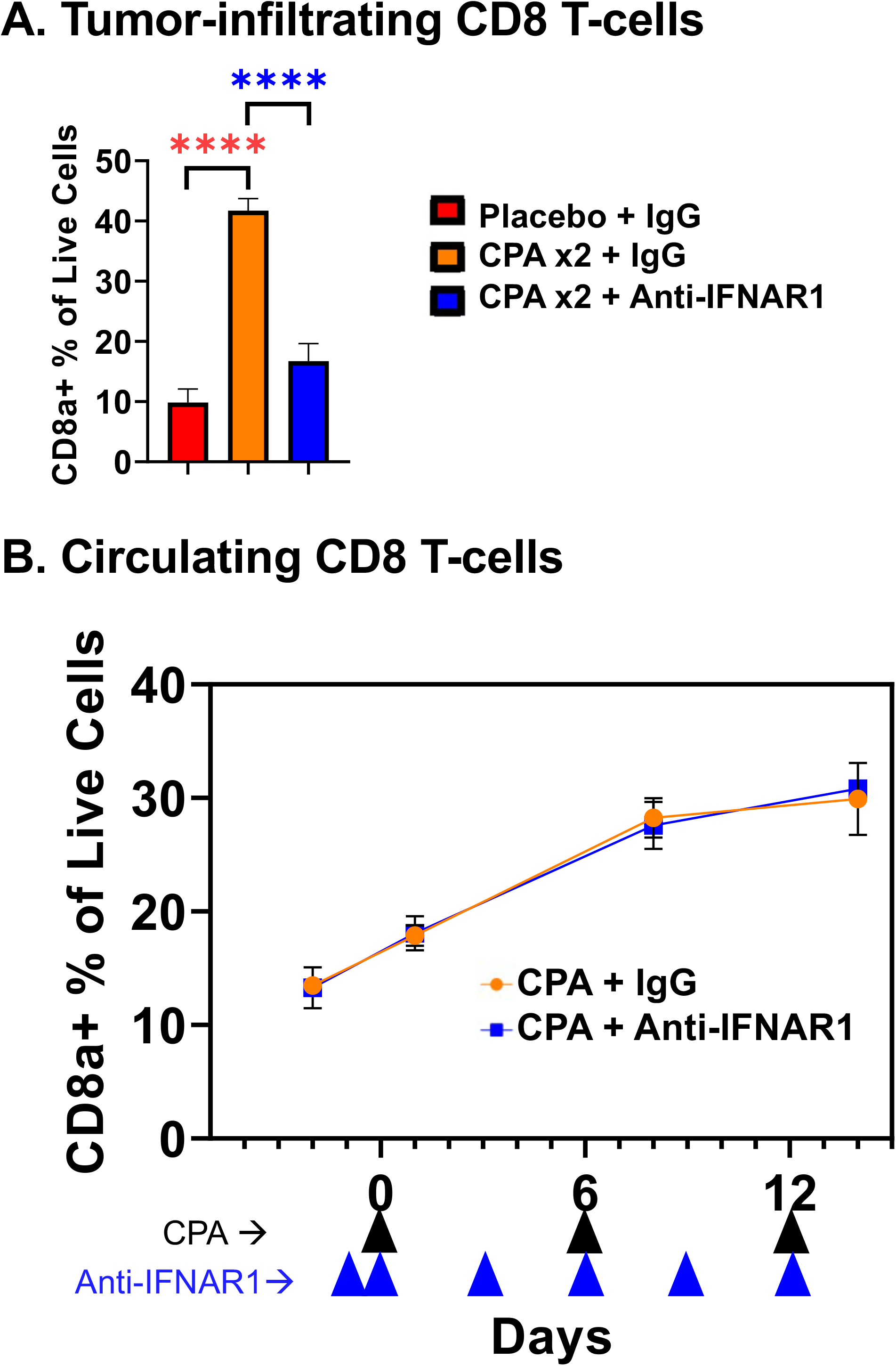
FACS analysis of blood and tumors from mice given metronomic CPA treatment with and without anti-IFNAR1 antibody. **A**. FACS analysis on treatment day 12 of tumor infiltrating CD8 T-cells from the mice shown in Fig. 8. Anti-IFNAR1 antibody treatment almost completely blocked CD8 T-cells from infiltrating the tumors. Data shown are mean +/-SEM values for n = 4 for placebo + IgG, n = 5 for CPA + IgG, and n = 6 for CPA + Anti-IFNAR1. **B**. FACS analysis of circulating CD8 T-cells from Fig. 8 mice. Circulating CD8 T-cells were not significantly different in the CPA + Anti-IFNAR1 group versus the CPA + IgG group. Data based on n = 3 for each group.

**Fig. S8.**
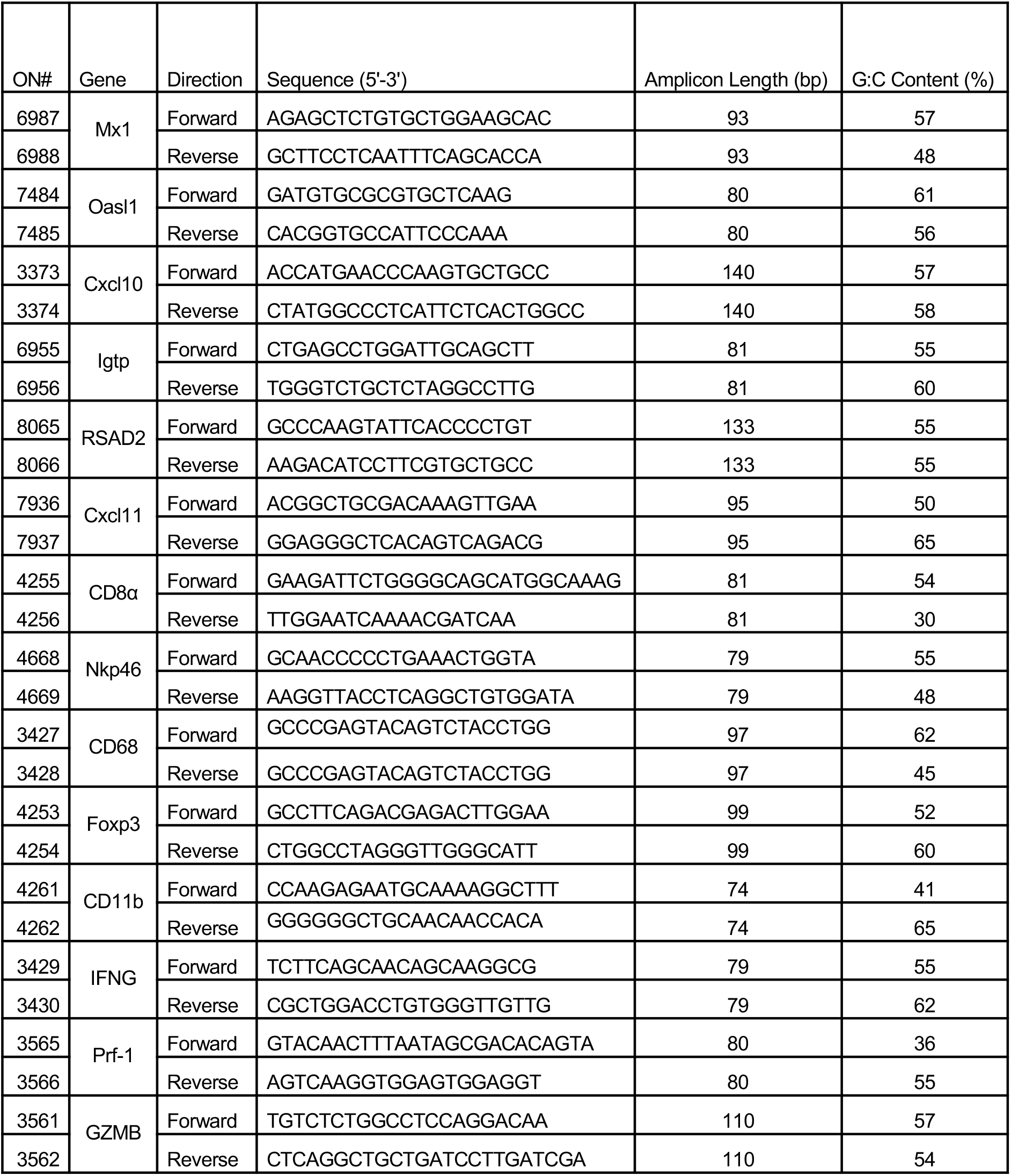
Gene-specific qPCR primer sequences, amplicon length and % GC content.

